# KSHV-encoded vIRF3 Cooperates with Cellular IRF4 to Drive Super-Enhancer Activity through Complex DNA Elements

**DOI:** 10.1101/2025.08.09.669493

**Authors:** Ziyan Liang, Haocong Katherine Ma, Christine B. Magdongon, Maureen E. Haynes, Jingyi Lu, Elizabeth T. Bartom, Eva Gottwein

**Affiliations:** Department of Microbiology-Immunology, Northwestern University, Feinberg School of Medicine, Chicago, IL 60611, USA; Departments of Biochemistry and Molecular Genetics and Preventive Medicine (Biostatistics and Informatics), Northwestern University, Feinberg School of Medicine, Chicago, IL 60611, USA

## Abstract

The Kaposi’s sarcoma-associated herpesvirus (KSHV) oncoprotein vIRF3 is essential for the survival of primary effusion lymphoma (PEL) cells. vIRF3 cooperates with cellular IRF4 to activate super-enhancers (SEs) driving oncogenes including MYC and IRF4 itself. However, the vIRF3/IRF4-responsive DNA sequences underlying this cooperation are unknown. Investigating the IRF4-SE, we mapped its vIRF3/IRF4-responsiveness to a complex ∼83 bp region, which retained cooperative activation by vIRF3 and IRF4 and was activated by vIRF3 but not IRF4 alone. vIRF3-mediated activation depended on an AP1 site, while the cooperation of vIRF3 with IRF4 required the DNA binding ability of IRF4 and IRF-related motifs that do not participate in canonical AP1-IRF (AICE) composite sites. These motifs are necessary but insufficient to confer responsiveness outside their native sequence context, suggesting that vIRF3/IRF4-mediated IRF4-SE activation requires an extended composite element. DNA pulldowns confirmed the importance of the identified motifs for association of vIRF3 and IRF4 with the IRF4-SE. A PEL MYC-SE similarly depended on an extended responsive element containing a critical AP1 site, within a functional AICE motif. Together, our results show that vIRF3 activates oncogenic SEs by co-opting complex genetic elements that may accommodate previously unknown IRF4 binding configurations, improving our understanding of vIRF3 and IRF4-dependent oncogenesis.

**Graphical Abstract:** 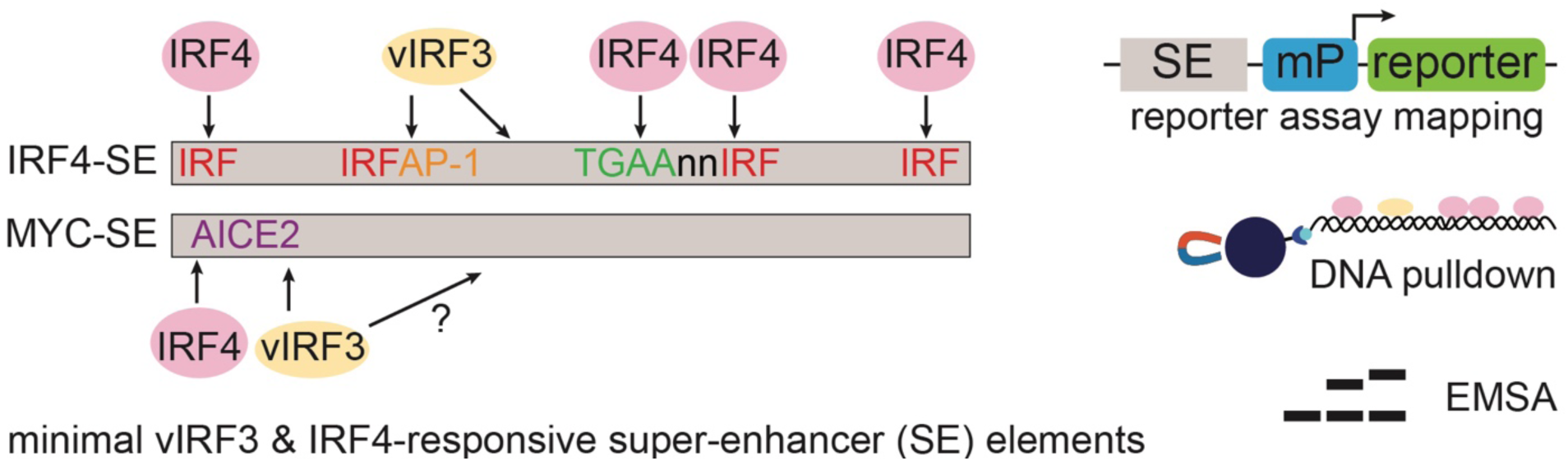

## Introduction

Kaposi’s sarcoma-associated herpesvirus (KSHV, also known as human herpesvirus 8) is the causative agent of primary effusion lymphoma (PEL) (1–3). The vast majority of PEL tumor cells exhibit the latent KSHV gene expression program. Notably, over 80% of PEL tumors are co-infected by Epstein-Barr virus (EBV). PEL tumor cells display the genetic rearrangements and gene expression features of post-germinal center B cells, including high expression of the hematopoietic lineage transcription factor (TF) interferon regulatory factor 4 (IRF4) (4–6). Accumulating evidence shows that the overexpression of IRF4 is critical for PEL pathogenesis. Specifically, IRF4 is essential for the survival and proliferation of all PEL-derived cell lines tested to date, irrespective of EBV co-infection (7–9). PEL shares this IRF4 dependency with several other hematologic malignancies, including multiple myeloma (10), the activated B cell-like subtype of diffuse large B cell lymphoma (11,12), anaplastic large cell lymphoma (13,14), and adult T-cell leukemia/ lymphoma (ATLL) (15). IRF4 dependency is finally observed in EBV-transformed lymphoblastoid cell lines (LCLs) (16,17), which result from the *in vitro* immortalization of primary B cells.

In PEL, other IRF4-dependent cancers, and LCLs, IRF4 occupies super-enhancers (SEs) (8,15,18–20). Super-enhancers are large clusters of enhancer elements, typically spanning >8 kb in contrast to <1kb regular enhancers (21,22). SEs drive exceptionally high transcription from promoters linked in 3D through looping factors. In cancer, SEs have been found to mediate the overexpression of essential proteins that promote cellular survival, proliferation, and other cancer-cell-specific features (23). One of the most critical SE-linked transcriptional targets of IRF4 in the IRF4-dependent cancers and LCLs is MYC. MYC itself is a TF with roles in proliferation, cell growth, and metabolism (24–26). The aberrant overexpression of MYC contributes to many cancers, due to genetic amplification or translocation, or transcriptional or post-transcriptional upregulation. IRF4-dependent cancers, including PEL, typically require the super-enhancer-mediated transcriptional activation of MYC. IRF4-dependent SEs furthermore drive the overexpression of the IRF4 gene itself and essential IRF4 cofactors (15,20–23), likely reinforcing IRF4 expression and function.

IRF4 is one of nine human interferon regulatory factor (IRF) TFs (27,28). IRFs share a highly conserved N-terminal DNA binding domain (DBD), followed by a variable linker domain, and a moderately conserved IRF-association domain (IAD) involved in interactions between IRFs and other protein partners. Several IRFs, including IRF4, additionally contain a C-terminal autoinhibitory region (AR). IRF DBDs interact with the core IRF DNA motif (5’-GAAA, referred to as IRF/GAAA here), which is found in tandem in the interferon-sensitive response element (ISRE; 5’-GAAAnnGAAA, where n is any nucleotide), for example, within the promoters of interferon-stimulated genes (ISGs). IRF4 expression is induced downstream of T and B cell receptor (TCR or BCR) signaling, such that the magnitude of IRF4 induction correlates with the strength and duration of receptor engagement (29–32). IRF4’s participation in TF complexes is governed by this variable expression level during immune cell differentiation and the availability of cofactors. IRF4 binds to the consensus IRF site more weakly than other IRFs, due to divergent features in its DNA-binding domain and autoinhibition through its AR domain (33). At low IRF4 expression levels, association with other TFs can relieve IRF4 autoinhibition and facilitate binding to composite sites. This paradigm of IRF4 activation is most established for the cooperation of IRF4 with ETS family members, PU.1 (encoded by *SPI1*) or SPIB, on ETS-IRF (EICE, 5’-GGAAnnGAAA) composite sites (33). Similarly, IRF4 and AP1 complexes, comprised of BATF and JUN subfamily members, bind to AP1-IRF composite elements (AICE, which have the consensus sequence 5’-TGAnTCA-n4-GAAA or GAAATGAnTCA, called AICE1 or AICE2, respectively) (34–36). IKZF1 (Ikaros) and IRF4 may finally recognize zinc finger-IRF composite sites (37). At the high concentrations of IRF4 expression that drive plasma cell differentiation, IRF4 homodimerizes on the ISRE, which is critical for the induction of the plasma cell master regulator PRDM1(also known as BLIMP1) and consequent plasma cell differentiation (30).

The exact identity of the SE elements and transcription factors that cooperate with IRF4 appear to differ between IRF4-dependent settings. Our published study used ChIP-seq to show that the latently expressed KSHV vIRF3 protein co-occupies several PEL super-enhancers with IRF4 and BATF. Expression of vIRF3 is required for the survival of PEL cell lines in culture (8,38,39). Published ChIP-seq, RNA-seq, Hi-ChIP, Western, and reporter data together support a model where vIRF3 and IRF4 positively regulate the PEL SEs linked to the *IRF4* and *MYC* genes, whose high expression is essential for the survival and proliferation of PEL cells (8,18,19). Targeting the vIRF3/IRF4-occupied SE-elements linked to IRF4 and MYC using CRISPR interference (CRISPRi) resulted in reduced expression of the linked genes and loss of the cultures (8,19). Similarly, knockdown of either IRF4 or vIRF3 resulted in reduced expression of genes linked to vIRF3/IRF4-occupied SEs (8,19), including IRF4 and MYC. While BATF is also essential in PEL cells, the reduction in viable PEL cell numbers following knockdown of BATF was delayed by several days compared to the more rapid loss of the culture after the inactivation of vIRF3 or IRF4, leaving the role of BATF less clear (8). The chromatin occupancy of JUN family members has not been studied in PEL cells and individual family members did not score as essential in CRISPR screens (8,9), suggesting potential redundancies for JUN family members as essential BATF binding partners.

Many questions remain regarding how vIRF3 cooperates with IRF4. vIRF3 shares homology with cellular IRFs in its N-terminal domain and C-terminal domains. While the N-terminal domain of vIRF3 is predicted to resemble the DBD of cellular IRFs structurally, it lacks the conserved amino acids that mediate the recognition of the IRF motif. Since vIRF3 is, therefore, not expected to bind to the IRF motif, it is unknown how vIRF3 is recruited to SEs. Our published ChIP-seq dataset identifies few high confidence vIRF3 peaks (8), including ∼60 that locate in SEs and overlap IRF4 peaks. HOMER motif analyses indicated significant enrichments of these peaks for composite sites, i.e., ISRE, AICE, and EICE sites, as well as IRF and AP1 sites. However, the actual TF occupancy and regulatory importance of these sites may depend on the cellular context, and it is unknown whether these motifs mediate vIRF3 and IRF4 function at the essential PEL SEs.

Here, we begin to address these questions by mapping the IRF4/vIRF3-responsive elements in the IRF4 and MYC-SEs. Our results suggest that vIRF3/IRF4-dependent SE activation in PEL is mediated by extended and complex composite genetic elements. Perhaps surprisingly, our data support a model for IRF4-and BATF-independent recruitment of vIRF3 to SE elements, which involves critical AP1 motifs in the IRF4-SE and one of two MYC-SEs. In contrast, the cooperation of vIRF3 with IRF4 depends on IRF-related motifs and the DNA binding ability of IRF4. Together, these data indicate that vIRF3 activates oncogenic SEs through complex genetic elements that may accommodate previously unknown IRF4 binding configurations.

## Material and Methods

### Biological Resources

293T/17 (“293T”) (ATCC, CRL-11268) were grown in Dulbecco’s Modified Eagle’s Medium (DMEM, Corning, 10-017-CV) containing 10% Serum Plus™ II Medium Supplement (Sigma-Aldrich 14009C-500ML, Batch Number: 21C421) and 10 μg/ml gentamicin (Gibco, 15710072). 293 (ATCC, CRL-1573) were grown in DMEM containing 10% fetal bovine serum (FBS, Corning, 35-010-CV) and 10 μg/ml gentamicin. BC-3 (ATCC, CRL-2277) were grown in RPMI 1640 (Corning, 10-040-CV), containing 20% FBS and 10μg/ml gentamicin. BJAB (obtained from Dr. Bryan Cullen at Duke University Medical Center) were grown in RPMI containing 10% FBS and 10μg/ml gentamicin. All cell lines were grown at 37°C with 5% CO2. pLS-mP (40) was a gift from Nadav Ahituv (Addgene plasmid # 81225; http://n2t.net/addgene:81225; RRID:Addgene_81225). The pLS-mP vector used here differs slightly from the vector we used previously (8). LVDP-CArG-RE-GPR was a gift from Stelios Andreadis (Addgene plasmid # 89762; http://n2t.net/addgene:89762; RRID:Addgene_89762). pGL4.72[hRlucCP] (Genbank Accession number AY738228) and pGL4.23[luc2/minP] (referred to as “pGL4.23” here, catalog number: E8411, GenBank Accession number DQ904455) were purchased from Promega, Madison, WI, USA. pLC-IRF4-IRES-Puro and related lentiviral vIRF3 and BATF expression vectors were described previously (8).

### Reagents and Specialized Equipment

All restriction enzymes, purchased in HF versions, where available, and the Q5 Site-Directed Mutagenesis Kit were from New England Biolabs (Ipswich, MA, USA). We used Taq polymerase (prepared as in 41) to amplify probes for DNA pulldown assays. Oligonucleotides, Ultramers, and gBlocks used for cloning were purchased from Integrated DNA Technologies (IDT, Coralville, IA, USA) and are listed in Table S1. Modified oligos used for DNA pulldown assays and EMSAs were also purchased from IDT and are listed in Tables S2 and S3. We performed Gibson Assembly (42) using an in-house protocol that is available on request. We used the Dual-Luciferase® Reporter Assay System (Promega, Madison, WI, USA, catalog number E1980) in combination with a VICTOR® Nivo™ Multimode Plate Reader with Dual Injectors from Perkin Elmer (Waltham, MA, USA). We imaged all gels and Westerns using a LI-COR (Lincoln, NE, USA) Odyssey-M Imaging System. Flow cytometry was performed with a BD FACSymphony™ A1 or FACSCanto™ II. Information for primary and secondary antibodies and control IgG is detailed in Table S4. Other reagents are listed in the relevant sections.

### Software

FACS data were analyzed in FlowJo v10 (BD).

### Statistical Analyses

We performed statistical analyses using GraphPad Prism 10 as detailed for each figure or panel. We performed two independent repeats for EMSAs and expression controls that included well-defined positive controls and had qualitatively reproducible results in these repeats. We performed at least three independent repeats for all figures that involved statistical analyses over replicates.

### Constructs and cloning procedures

All newly cloned vectors were verified by Sanger Sequencing (ACGT, Wheeling, IL, USA) or, where the NEB Q5 Site-Directed Mutagenesis Kit was used, by full plasmid sequencing (Plasmidsaurus, South San Francisco, CA, USA).

To clone pGL4.72-RSV-LTR, the RSV-LTR was PCR-amplified from a lentiviral vector using primers 393 and 394 and cloned between the BglII and HindIII sites of pGL4.72[hRlucCP] using T4 DNA ligase.

The following constructs were cloned by Gibson Assembly of the relevant PCR products with XhoI/HindIII digested pGL4.23[luc2/minP]. For pGL4.23-IRF4-SE^500bp^ we inserted a PCR fragment we amplified from the published vector pLS-IRF4-SE-eGFP (8) using primers 4850/4851; for pGL4.23-IRF4-SE^69-151^, the relevant fragment was amplified from pGL4.23-IRF4-SE^500bp^ using primers 5056/5057; for pGL4.23-MYC-SE^500kb^ a 327bp fragment was inserted, which was originally amplified from genomic DNA and re-amplified using primers 4959/4960; for pGL4.23-MYC-SE^375kb^ a 299 bp fragment was amplified from genomic DNA using primers 4854/4855. PCR products for pGL4.23-IRF4-SE^500bp^-50 bp and 18 to 22 bp deletion mutants were generated using primers 4957 and 5048 (Δ1-50), 4850 and 4958 (Δ451-500), or by overlap PCR using outer primers 5048/5049 and two PCR fragments as templates, which were in turn amplified using forward primer 5049 and mutant-specific reverse primers or mutant-specific forward primers and reverse primer 5048.

The pGL4.23-IRF4-SE^69-151^-GAAA1/2/3/4 mutants, -AICE-like mutant, -ISRE-like mutant, and the pGL4.23-IRF4-SE^500bp^-Δ51-151 mutant were cloned using the NEB Q5 Site-Directed Mutagenesis Kit using pGL4.23-IRF4-SE^69-151^ or pGL4.23-IRF4-SE^500bp^ as templates. pGL4.23-IRF4-SE^500bp^-Δ51-151 was not used for reporter assays in this paper but served as a template for pLS-mP cloning. To clone the pGL4.23-IRF4-SE^69-151^-4GAAA, TGAnTCA, and TGAA mutants, we inserted synthesized dsDNA fragments (gBlocks) 5451, 5454, or 5455, respectively, into the XhoI/HindIII-digested pGL4.23 vectors using Gibson Assembly. To clone pGL4.23-IRF4-SE^69-151^-R1^old^ and -R2, we performed Gibson Assembly of the XhoI-HindIII-digested vector with gBlocks (IDT) 5456 and 5457, respectively. pGL4.23-IRF4-SE^69-151^-R1^old^ was not used for reporter assays in this paper but served as a template for cloning below. pGL4.23-IRF4-SE^69-151^-R1, which retains the A between the AP-1 and 2^nd^ GAAA motif, was cloned using the Q5 Site-Directed Mutagenesis Kit, using primers 5618/5619, and pGL4.23-IRF4-SE^69-151^-R1^old^ as template. pGL4.23-IRF4-SE-87-107, 87-129, 87-129 and pGL4.23-R3/R4/R5 were cloned by joining annealed oligos or Ultramers with XhoI/HindIII-digested pGL4.23 vector using T4 DNA ligase.

pGL4.23-MYC-SE^500kb^ 40 bp deletion mutants were generated by overlap PCR using primers 5048/5049 and two PCR fragments as templates, which were in turn amplified using forward primer 5049 and mutant-specific reverse primers or mutant-specific forward primers and reverse primer 5048. Mutant overlap PCR products were inserted into the XhoI/HindIII-digested pGL4.23 vector using Gibson Assembly. pGL4.23-MYC-SE^500kbp^-AICE2, TGAnTCA, and GAAA mutants and the pGL4.23-MYC-SE^375kbp^ TGAnTCA mutant were cloned using the NEB Q5 Site-Directed Mutagenesis Kit and the relevant wt vector as template.

To clone the pLS-IRF4-SE^500bp^ and -IRF4-SE^69-151^ reporters, we used primers 5620/5621 and 5622/5623 to amplify the relevant sequences from pGL4.23-IRF4-SE^500bp^ and pGL4.23-IRF4-SE^69-151^, respectively. The products were cloned into XbaI-digested pLS-mP using Gibson Assembly. pLS-mP-IRF4-SE^500bp^-Δ69-151 was cloned by inserting two overlapping PCR fragments amplified from pLS-mP-IRF4-IRF4-SE^500bp^ with primers 5620/5523 and 5524/5621 into XhoI/XbaI-digested pLS-mP-IRF4-SE^500bp^ using Gibson Assembly. To clone the remaining pLS-mP-IRF4-SE^69-151^ mutants, we used primers specified in Table S1 to amplify the relevant fragments from the corresponding pGL4.23 constructs. The products were inserted into XhoI/XbaI-digested pLS-mP-IRF4-SE^500bp^ by Gibson Assembly.

LVDP-CArG-RE-GPR was cut using AgeI and EcoRI. Fragments containing the minimal promoter, minP, and minP-IRF4-SE^500bp^ were amplified with primers 5982/5983 and 5982/5984 from template pLS-mP-IRF4-SE^500bp^ and inserted into the LVDP-CArG-RE-GPR backbone by Gibson Assembly, resulting in plasmids LVDP-minP and LVDP-minP-IRF4-SE^500bp^, which were only used for further cloning here. To replace the original LVDP-CArG-RE-GPR dsRed-Express 2 cassette with mCherry, we amplified an mCherry-containing fragment with primers 6000/6001 and an IRF4-SE^500bp^-ZsGreen fragment from LVDP-minP-IRF4-SE^500bp^ with primers 6002/6003. We obtained the hPGK-cHS4 fragment by restriction digest of LVDP-minP-IRF4-SE^500bp^ with NheI and XbaI. The three resulting fragments were inserted into XhoI/EcoRI-digested LVDP-minP-IRF4-SE^500bp^ using Gibson Assembly to generate mLVDP-minP-IRF4-SE^500bp^. We generated a minP control vector, mLVDP-minP, by excising the minP fragment from LVDP-minP using AgeI and EcoRI and replacing the corresponding fragment of mLVDP-minP-IRF4-SE^500bp^ using T4 DNA ligase. mLVDP-minP-IRF4-SE^500bp^ and mLVDP-minP were only used for cloning in this study. To insert a puromycin resistance cassette downstream of mCherry, we amplified a P2A-Puro fragment from pL2M-MPH-2A-Puro (43) with primers 6058/6059, and mCherry and hPGK fragments from mLVDP-minP-IRF4-SE500 (with primers 6060/6061 and 6062/6063, respectively). The resulting three fragments were inserted into BamHI-digested mLVDP-minP-IRF4-SE^500bp^ to generate pLVDP-mCherry-P2A-Puro-IRF4-SE^500bp^. pLVDP-mCherry-P2A-Puro-IRF4-SE^500bp^-Δ69-151 and -IRF4-SE^69-151^ were cloned by excising the relevant fragments from the corresponding pLS-mP vectors using EcoRI and XbaI and inserting them into EcoRI- and XbaI-digested pLVDP-mCherry-P2A-Puro using T4 DNA ligase. To clone pLVDP-mCherry-P2A-Puro-minP, we excised the relevant fragment from mLVDP-minP using PacI and EcoRI and replaced the corresponding fragment of pLVDP-mCherry-P2A-Puro-IRF4-SE^500bp^ using T4 DNA ligation. To generate pLVDP-mCherry-P2A-Puro-IRF4-SE^200bp^, we amplified the first 200 bp of IRF4-SE^500bp^ from pGL4.23-IRF4-SE^500bp^ using primers 6003/6091 and used Gibson Assembly to insert the resulting fragment into EcoRI/XbaI-digested pLVDP-mCherry-P2A-Puro-IRF4-SE^500bp^. To generate pLVDP-mCherry-P2A-Puro-IRF4-SE^500bp^-AP1mut, we amplified AP1 mutant IRF4-SE^500bp^ fragments using primers 6092/6093 and 6094/6095 and used Gibson Assembly to insert them into EcoRI/XbaI-digested pLVDP-mCherry-P2A-Puro-IRF4-SE^500bp^. pLVDP-mCherry-P2A-Puro-MYC-SE^500kb^ and an AP-1 mutant version were cloned by inserting PCR fragments generated using primers 5086/6096 and wt or AP-1 mutant templates pGL4.23-MYC-SE^500kb^ into EcoRI/XbaI-digested pLVDP-mCherry-P2A-Puro-IRF4-SE^500bp^ using Gibson Assembly. To generate pLVDP-mCherry-P2A-Puro-MYC-SE^500kb^/^81-247bp^, we amplified the relevant fragment from pGL4.23-MYC-SE^500kb^ using primers 6097 and 6098 and performed Gibson Assembly with EcoRI/XbaI-digested pLVDP-mCherry-P2A-Puro-IRF4-SE^500bp^. To generate AP-1 mutant pLVDP-mCherry-P2A-Puro-MYC-SE^500kb^/^81-247bp^, we amplified the relevant mutant fragments using primers 6097/6099 and 6100/6098 and cloned them into EcoRI/XbaI-digested EcoRI/XbaI-digested pLVDP-mCherry-P2A-Puro-IRF4-SE^500bp^ using Gibson Assembly.

We cloned pLC-EBFP2-IRES-Puro, pLC-vIRF3-IRES-Puro, and pLC-BATF-IRES-Puro by moving the relevant expression cassettes between published vectors (8), using AgeI and BamHI and T4 DNA ligase. To clone pLC-BATF2-IRES-Puro and pLC-BATF3-IRES-Puro, which express untagged BATF2 and BATF3, respectively, we ordered gBlocks 6182 and 6183 containing HA-tagged expression cassettes, amplified the related untagged expression cassettes using primer 6184/6185 or 6184/6186, and joined them with AgeI- and BamHI-digested pLC-IRES-Puro using Gibson Assembly. pLC-JUNB-IRES-Puro was cloned by inserting a PCR fragment amplified from genomic DNA using primers 5481 and 5482 into AgeI/BamHI-digested pLC-IRES-Puro backbone, using Gibson assembly. JUNB coding sequences were confirmed to match the reference sequence by Sanger sequencing. To clone pLC-IRF4^R98A/C99A^-IRES-Puro, two mutant fragments were amplified using primers 4087/5479 and 5480/4764 and pLC-IRF4-IRES-Puro as template and inserted into the AgeI/BamHI-digested pLC-IRES-Puro backbone using Gibson Assembly. Sanger Sequencing confirmed sequences with primers 4259 and 2154.

### Dual-Luciferase reporter assay

293 cells were plated at 1 x 10^5^ cells per well of a 24-well plate one day before transfection. Cells were co-transfected with 50 ng of pGL4.23 Firefly luciferase reporter plasmid, 50 ng of an RSV-LTR-driven Renilla luciferase internal control plasmid pGL4.72-RSV-LTR, and 200 ng pLC-IRES-Puro-based expression plasmids, i.e., 100 ng of pLC-vIRF3-IRES-Pur and/or pLC-IRF4-IRES-Puro, and/or 100 or 200 ng pLC-EBFP2-IRES-Puro. For reporter assays including BATF/JUNB, we also used 100 ng per vector and the total amount of pLC-IRES-Puro plasmids was adjusted to 400 ng with pLC-EBFP2-IRES-Puro as filler. A Master Mix of 50 μl Opti-MEM and Lipofectamine LTX (Invitrogen # 15338100) [DNA(μg): LTX (μl)=1:3] per reaction was prepared and incubated at room temperature for 5 min. Then the master mix was mixed with plasmid combinations described above and incubated at room temperature for 15 min before addition into the wells. Approximately 24 hours post-transfection, media were aspirated, and cells were lysed using 100 μl of 1X Passive Cell Lysis Buffer (Promega) per well for 15 min. 5 μl supernatant of each lysate was transferred to a 96-well half-well white plate and Dual-Luciferase reporter assays were performed using the Dual-Luciferase® Reporter Assay System (Promega) according to manufacturer specifications on a Victor Nivo luminescence and fluorescence plate reader with dual injectors.

### Lentivirus production and concentration

Lentivirus production was performed as described. Briefly, 293T/17 were plated at 12.5 x 10^6^ cells per 15 cm dish or 8 x 10^5^ per well of a 6-well plate one day prior to transfection. Lentiviral transfer plasmids, psPAX2, and pMD2.G were co-transfected at 45:35:25 molar ratios using the polyethyleneimine (PEI, Polysciences) transfection method. The medium was changed 4∼6 hours after transfection. Three days after transfection, the supernatants containing lentiviral particles were harvested and filtered through 0.45 μm-pore-size filters to remove cell debris. pLS-mP-based LVs were concentrated using Amicon Ultra centrifugal filter units (Millipore UFC910024). The concentrated LV preparations were stored in aliquots at −80°C for future use. Filtered pLVDP-containing media were directly stored in aliquots at −80°C without concentration.

### RNA-based and Functional Titration of Lentiviruses

For pLS-mP lentiviruses, we performed both RNA-based and functional titration. For RNA-based titration, lentiviral RNA was purified from supernatants containing viruses using the NucleoSpin® RNA Virus kit (Takara #740956.50). The viral RNA was eluted in 50 μl RNase-free water provided with the kit and subjected to DNase treatment using the RQ1 DNase kit (Promega, #M6101), following the manufacturer’s protocol. Lentiviral genomic RNA copy numbers were quantified by reverse transcription quantitative PCR (RT-qPCR) using the Lenti-X qRT-PCR titration kit (Takara, #631235) following the manufacturer’s instructions, including no template controls (NTC, i.e., dilution buffer or RNase-free water), and no reverse transcriptase controls. A standard curve of threshold cycle (Ct) values versus copy numbers (log scale) was generated based on serial dilutions of the Lenti-X RNA Control template and used to calculate lentiviral copy numbers determined by RT-qPCR of serial dilutions of each viral RNA sample. In addition to RNA-based titration, we titered each lentivirus functionally using flow cytometry. We plated BC-3 in 12-well plates at a density of 4 x 10^5^ cells/ml and transduced cells with serial dilutions of viral supernatants in the presence of 5 μg/ml polybrene. One day after transduction, the medium was replaced, and cells were split to 3 x 10^5^/ml. Three days post-transduction, the percentage of GFP-expressing cells at each dilution was assessed by flow cytometry and functional titers (transducing units per milliliter) were calculated from dilutions generating single integration events, i.e., yielding <30% of GFP-positive cells. The relative RNA-based titers and functional titers were largely consistent across samples and, therefore, the functional titers were used to calculate the appropriate volume of viral supernatants for subsequent experiments. For pLVDP viruses, we only performed functional titrations in BC-3 cells as described above, calculating the functional titer from the percentage of mCherry-expressing cells two days after transduction.

### Lentiviral-based reporter assays in PEL cell lines

For pLS-mP viruses, BC-3 cells were plated at 4 x 10^5^ cells/ml in 12-well plates with 5 μg/ml polybrene. The lentiviruses containing SE-element-driven EGFP reporter were transduced at MOI of ∼1. The medium was changed one day after transduction. Three days after transduction, cells were subjected to flow cytometry and EGFP mean fluorescence intensity (MFI) was determined in EGFP-positive cells.

The pLVDP dual-reporter lentiviruses were transduced at MOI of 0.3 into BC-3 cells as described above. One day after transduction, we changed to medium with 1.2∼1.5 μg/ml Puromycin to select transduced cells. Three days after transduction, cells were subjected to flow cytometry, gated for live, single, mCherry-positive cells and reported ZsGreen mean fluorescence intensity in this population.

### Western blot

293 cells were transfected as described above with protein-expressing vector in 6-well plate, scaling by surface area. Cells were harvested one day after transfection. 293 cells were washed twice with PBS. The adherent cells were lysed by 100∼200 μl RIPA buffer (50 mM Tris pH8.0, 150 mM NaCl, 1% IGEPAL CA-630, 0.5% sodium deoxycholate, 0.1% SDS) supplemented with protease inhibitors and then scraped and transferred into precooled 1.5 ml tubes. The lysates were incubated on ice for 20∼30 min and sonicated for 6 cycles (30 sec on, 30 sec off). Insoluble debris was removed by centrifugation at 16,000 g for 20 min. The supernatants were transferred to a fresh tube and quantified using a BCA assay. BC-3 cells were subjected to the same procedure, except that BC-3 cells were centrifuged to remove the supernatant and washed with PBS. For whole cell lysates, 10 μg proteins were resolved in Bis-Tris gels and then transferred to 0.45 μm nitrocellulose membrane. Except for BATF, membranes were blocked using TBS (20 mM Tris, 137 mM NaCl, pH 7.6) containing 5% w/v non-fat milk powder. Next, membranes were incubated with primary antibody in TBST (TBS containing 0.1% Tween-20)-5% w/v non-fat milk powder (0.1% Tween-20). Membranes were washed three times with TBST for 15 min each time before incubating with IDye-800-conjugated secondary antibody in TBST with 5% milk. Membranes were washed three times with TBST for 15 min each time and then imaged on a LI-COR Odyssey-M Imaging System. For BATF, we used the same protocol, but blocked membranes using LI-COR Intercept (TBS) Blocking Buffer and Intercept® T20 (TBS) Antibody Diluent.

### DNA pull-down assay

The 5’-biotinylated primers listed in Table S1 were ordered from IDT. We performed multiple PCR reactions using the above pGL4.23 vectors as template and in house Taq polymerase. PCR products were precipitated using 1/10 volume of 3 M Sodium Acetate, 50 μg/ml GlycoBlue, and 3 volumes of 100% ethanol. The mixture was incubated at −20°C for at least 1 hour and then subjected to centrifugation for 30min at 4°C and 16,000 g. The pellet was washed twice with ice-cold 70% ethanol, air dried, and resuspended in nuclease-free water. The products were then agarose gel purified (IBI Scientific Gel Extraction & PCR Cleanup Kit) and eluted in nuclease-free water. The concentration of the probes was determined using a Qubit Fluorometer.

To prepare nuclear extracts (NEs) used in EMSAs, 1 x 10^7^ BC-3 cells were centrifuged and washed twice with cold PBS. The cell pellet was resuspended with 400 μl ice-cold Hypotonic buffer [10 mM HEPES-KOH, pH 7.9, 10 mM KCl, 0.1 mM EGTA, 1.5 mM MgCl2, 1 mM DTT, 0.5 mM PMSF] and incubated on ice for 10 min. After adding 25 μl 10% Igepal CA-630, the sample was subjected to a quick spin to pellet the nuclear fraction. The nuclear pellet was resuspended with 40 μl Nuclear Extract buffer (20 mM HEPES-KOH pH 7.9, 420 mM NaCl, 0.2 mM EDTA, 1.5 mM MgCl2, 25% glycerol, 0.5 mM DTT, 0.5m M PMSF, and protease inhibitor cocktail) and incubated on ice for 30 min with brief vortexing every 10 min. The sample was centrifuged at maximal speed for 12 min and the supernatants were collected and proceeded to BCA assays for protein quantification.

50 μg of BC-3 NEs were incubated with 500 ng of biotinylated DNA probes overnight at 4°C in the pull-down buffer (10 mM HEPES-KOH pH7.9, 50 mM KCl, protease inhibitors). Poly[d(I-C)] was added to a final concentration of 10 ng/μl to reduce non-specific binding. 20 μl of Dynabeads M280 Streptavidin (Invitrogen #11205D) were pre-washed three times with washing buffer (10 mM HEPES-KOH pH 7.9, 50 mM KCl, 0.05% Tween-20) and resuspended with 40 μl washing buffer. The washed beads were added to the protein-probe mix and incubated for 2 hours at 4°C. The beads were then washed six times with the washing buffer and resuspended in 40 μl resuspension buffer (10 mM HEPES-KOH pH 7.9, 50 mM KCl). The resulting samples were subjected to Western blot following the description above but using running buffer containing NuPAGE™ Antioxidant (1:400, Invitrogen NP0005) in the upper chamber. DNA pull-down assay using 293T NE were performed similarly, using either 25 μg or 50 μg NE, with similar relative results.

### Nuclear Co-Immunoprecipitation (Co-IP) assay

3 x 10^6^ 293T cells were plated in 10 cm dish one day before transfection. 6 μg of each protein expression plasmid was transfected using the PEI method and the EBFP2 plasmid was used to to adjust the total amount of DNA to 12 μg/dish. Two days after transfection, nuclear extracts were prepared using the Active Motif (Carlsbad, CA, USA) Nuclear Extract Kit (#40010) according to instructions. Protein concentrations were quantified using BCA assay (Pierce).

Co-IP was performed with reagents provided with the Active Motif Nuclear Complex Co-IP kit (#54001). Specifically, 100 μg of nuclear extracts were incubated with 1.5 μl anti-IRF4 (Cell Signaling, #D9P5H, rabbit) in IP incubation buffer overnight at 4°C. 20 μl of Dynabeads protein A magnetic beads (ThermoFisher, #10002D) were the next day and the mixture was incubated for another 2 hours at 4°C. The beads were then washed six times and resuspended in IP wash buffer supplied with the kit. The Co-IP samples together with nuclear extracts were analyzed by Western Blot, using rat anti-IRF4 (ThermoFisher, clone 3E4, catalog 14-9858-82) as primary antibody to detect IRF4.

### Electrophoretic Mobility Shift Assay (EMSA)

5’ IRDye® 700-labeled oligonucleotides were purchased from IDT. The oligos were annealed in annealing buffer (10 mM Tris (pH 7.5∼8), 50 mM NaCl, 1 mM EDTA) in thermocycler following the procedure (95°C for 3 min and 37°C for 1 hour). The resulting probes were diluted to 50nM as working concentration. 3 x 10^6^ 293T cells were plated in 10 cm dish one day before transfection. 3 μg of each protein-expressing plasmid was transfected using PEI method, using the EBFP2 plasmids to adjust total amount of DNA to 9 μg/dish. 2 days after transfection, nuclear extracts were harvested using a nuclear extract kit (Active Motif #40010) following the manual. Protein concentrations were quantified using BCA assay (Pierce). EMSA was performed with reagents included in the Odyssey EMSA kit (LI-COR, #82907910). Pre-mixed reagents [10x binding buffer, 25 mM DTT/2.5% Tween-20, 1 μg/μl Poly (dI-dC)], 5μg of nuclear extracts and probes were incubated on ice for 20 mins. When performing supershifts, antibodies were added subsequently, and reactions were incubated for another 20 mins on ice. Orange loading dye supplied in the kit was added and samples were separated on DNA retardation gels (Invitrogen™, EC6365BOX) using 1x TBE as the running buffer (Fisher, BP1333-1). Gels were imaged by LI-COR Odyssey-M Imaging System.

## Results

### vIRF3/IRF4-responsive IRF4-SE sequences map to a complex 83 bp DNA element

We have previously used lentiviral EGFP-based reporter assays in 293 cells to demonstrate that vIRF3 and IRF4 positively regulate a 500 bp sequence centered on the overlapping major vIRF3 and IRF4 ChIP-seq peaks in the PEL IRF4-SE, located ∼63 kb upstream of the IRF4 transcription start site (8). However, vIRF3/IRF4-mediated reporter activation was relatively weak (∼2 fold), and the requirement for vector titration makes the transduction-based assay cumbersome to scale for mutagenesis studies. Transfection-based luciferase reporter assays recapitulate results from integrated lentiviral reporters but can offer a higher dynamic range (44). To facilitate mapping the sequence requirements for vIRF3/IRF4-mediated IRF4-SE activation, we first tested whether transfection-based dual-luciferase reporter assays recapitulate our published result. We inserted the published 500bp IRF4-SE element (Fig. S1, IRF4-SE^500bp^) fragment upstream of the minimal promoter (min. P) in the firefly luciferase (FLuc) reporter plasmid pGL4.23 (Promega). We then co-transfected the resulting reporter vector with an internal Rous sarcoma virus long terminal repeat (RSV-LTR) driven *Renilla* luciferase (RLuc) control vector (pGL4.72/RSV-LTR) and combinations of vIRF3, IRF4, and EBFP2-expressing vectors into 293 cells (Fig. 1A, B). None of the tested combinations affected RLuc expression from the RSV-LTR internal control reporter (Fig. S2A). While IRF4 or vIRF3 alone did not significantly activate relative IRF4-SE^500bp^ FLuc reporter expression (Fig. 1C), IRF4-SE reporter expression was ∼15-fold induced after co-transfection of vIRF3 and IRF4 (Fig. 1C, S2B). These results reproduce our published data, with greatly improved sensitivity, pointing to cooperative activation of this IRF4-SE element by vIRF3 and IRF4.

**Figure 1.**
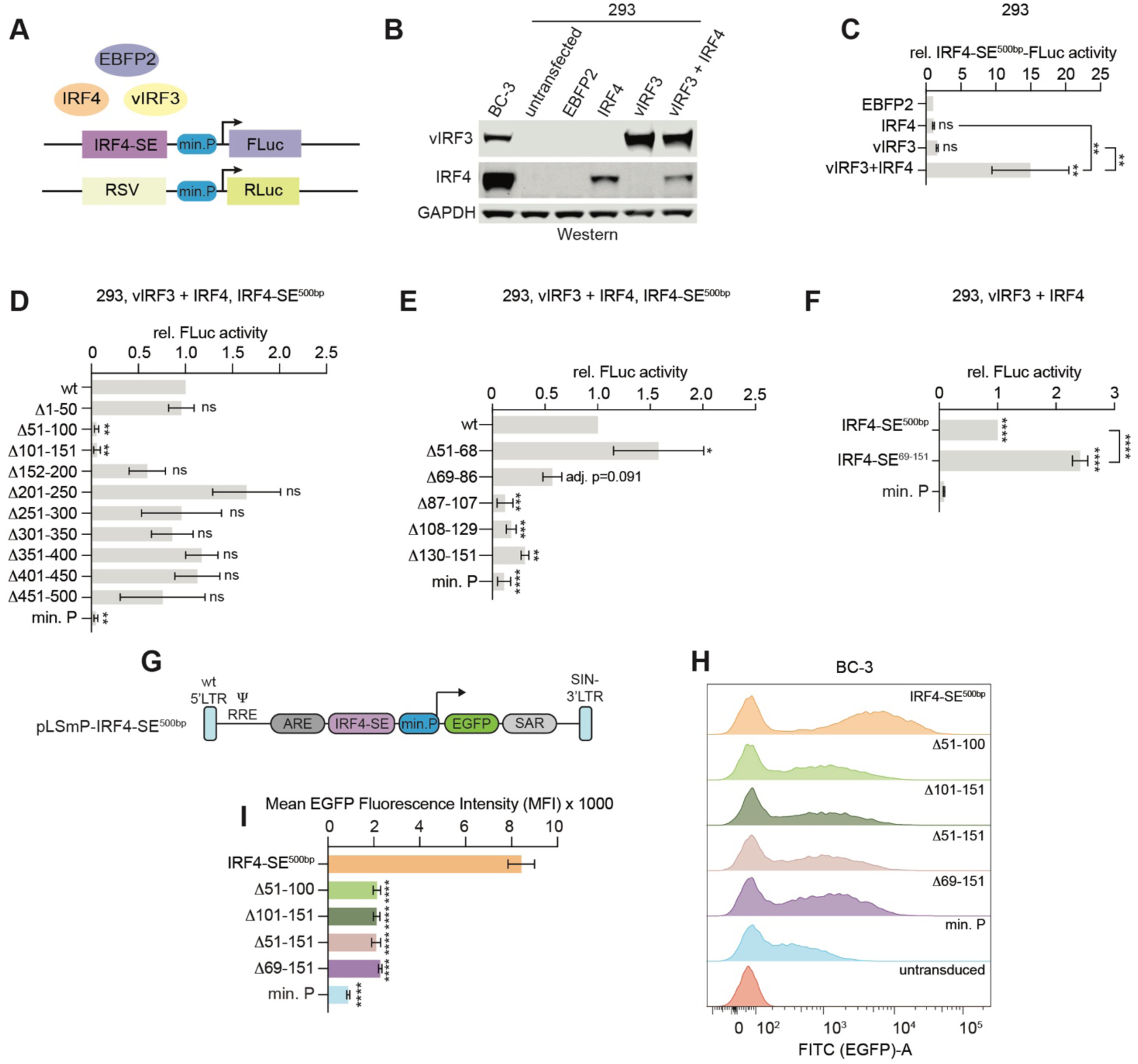
The vIRF3/IRF4-responsive element within the IRF4-SE maps to an 83bp region. **(A)** Schematic of the approach used for dual-luciferase reporter assays. See text for abbreviations. **(B)** Western blot analyses of IRF4 and vIRF3 expression one day after transfection of the indicated expression vectors into 293, under the conditions used for reporter assays. GAPDH served as a loading control. BC-3 served as a control for expression levels in PEL cells. Representative of 2 independent repeats. **(C)** Results from dual-luciferase reporter assays as described in panel (A). We sequentially normalized results from FLuc reporters to those from the RSV-RLuc internal control and then results from TF combinations to those from EBFP2, which we set to 1. Significance is relative to EBFP2 unless indicated by lines. **(D)** Results from dual-luciferase reporter assays as in panel (C), but co-transfecting wt or 50bp IRF4-SE^500bp^ deletion mutant reporters with EBFP2, or vIRF3 and IRF4. For each reporter, results from vIRF3/IRF4-cotransfected samples were normalized to those from EBFP2, as in panel (C), and additionally to the result from the wt IRF4-SE^500bp^ reporter. min. P: minimal promoter-only Fluc control vector. Significance is indicated relative to the wt vector. **(E)** Results from dual-luciferase reporter assays as described in panel (D), measuring the effect of 18-20 bp deletions between nts 51-151 of the IRF4-SE^500bp^. Significance is indicated relative to the wt vector. The reduction of Δ69-86 activity is significant when compared to IRF4-SE^500bp^ using a one-way ANOVA followed by Dunnett’s post hoc tests (*adj. p* = 0.037) and we therefore included this region in panels F-G. **(F)** Results from dual-luciferase reporter assays as described in panel (D) but comparing the IRF4-SE^500bp^ reporter and a reporter containing nts 69-151 of IRF4-SE^500bp^ (IRF4-SE^69-151^). Significance is relative to min. P unless indicated by lines. **(G)** Schematic of the pLS-mP (40) lentiviral reporter vector used in this figure, showing wild-type (wt) 5’ long terminal repeats (5’LTR), packaging signal (4′), Rev response element (RRE), antirepressor element #40 (ARE), a scaffold-attached region (SAR), IRF4-SE element, minimal promoter (min. P), EGFP coding sequence, and self-inactivating (SIN) 3’LTR. **(H)** Representative histogram comparing lentiviral EGFP reporter expression 3 days after transduction of BC-3 cells with reporters containing the wt IRF4-SE^500bp^ sequence or the indicated mutants at equal MOI (∼1). **(I)** Quantification of EGFP mean fluorescence intensity of EGFP-positive BC-3, defined based on an untransduced negative control, over 3 independent experiments as in panel (H). Significance is relative to the wt IRF4-SE^500bp^ reporter. Throughout, error bars represent SD from 3 biological replicates for bar graphs. ns, not significant; *, *adj. p <* 0.05; **, *adj. p <* 0.01; ***, *adj. p <* 0.001; ****, *adj. p <* 0.0001, calculated using One-Way ANOVA followed by Tukey’s post hoc tests.

To begin mapping the sequences that confer vIRF3/IRF4-dependent IRF4-SE^500bp^ reporter activation, we initially introduced ∼50bp deletions. We co-transfected resulting mutants with expression vectors for the EBFP2 negative control or vIRF3 and IRF4. IRF4-SE^500bp^ mutants lacking either nucleotides (nts) 51-100 or 101-151 showed strongly reduced reporter activity (Fig. 1D). In contrast, other deletions had a minimal or no impact, suggesting that nts 51-151 harbor the vIRF3/IRF4-responsive element(s). Smaller 18-22 bp deletions within this region, except for Δ51-68bp, impaired vIRF3/IRF4-mediated reporter activation (Fig. 1E), suggesting that a relatively large ∼83bp region mediates vIRF3/IRF4-mediated reporter activation. This fragment (IRF4-SE^69-151^) was sufficient for full reporter responsiveness to vIRF3/IRF4 in 293 cells (Fig. 1F) and, therefore, represents a minimal vIRF3/IRF4-responsive element within the IRF4-SE.

To test whether the identified minimal region is similarly critical for IRF4-SE activity in PEL cell lines, we transduced the PEL cell line BC-3 with lentiviral EGFP-based enhancer element reporters containing the wildtype (wt) IRF4-SE^500bp^ fragment or Δ51-100, Δ101-151, Δ51-151 or Δ69-151 mutants from above (Fig. 1G-I). Each deletion strongly reduced IRF4-SE^500bp^ reporter expression, confirming the regulatory importance of the minimal element in PEL cell lines. Thus, results from both 293 and the PEL cells point to the critical importance of a ∼83bp minimal vIRF3/IRF4 responsive element within the IRF4 SE, and we therefore focused our subsequent experiments on this region.

### The minimal vIRF3/IRF4-responsive IRF4-SE element lacks a functional AICE motif

Inspection of the identified minimal vIRF3/IRF4 responsive element within the IRF4-SE revealed four IRF/GAAA motifs (Fig. 2A), including one within a candidate ISRE-like motif (<u>TG</u>AAnnGAAA, underlined nucleotides diverge from the consensus). IRF4 tolerates one and even two-nucleotide variations from the IRF/GAAA consensus in the context of composite sites (30,34), although the sequence above would likely have considerably lower affinity to IRF4 than a consensus ISRE. The second IRF motif is close to a canonical AP1 motif (TGAnTCA), but not within the 4 nt spacing that is typical for an AICE1 site, which has the consensus sequence TGAnTCA-n4-GAAA. AICE and ISRE motifs are binding sites for IRF4/BATF/JUN complexes and IRF4 homodimers, respectively. In contrast, isolated IRF motifs are not thought to recruit IRF4.

**Figure 2.**
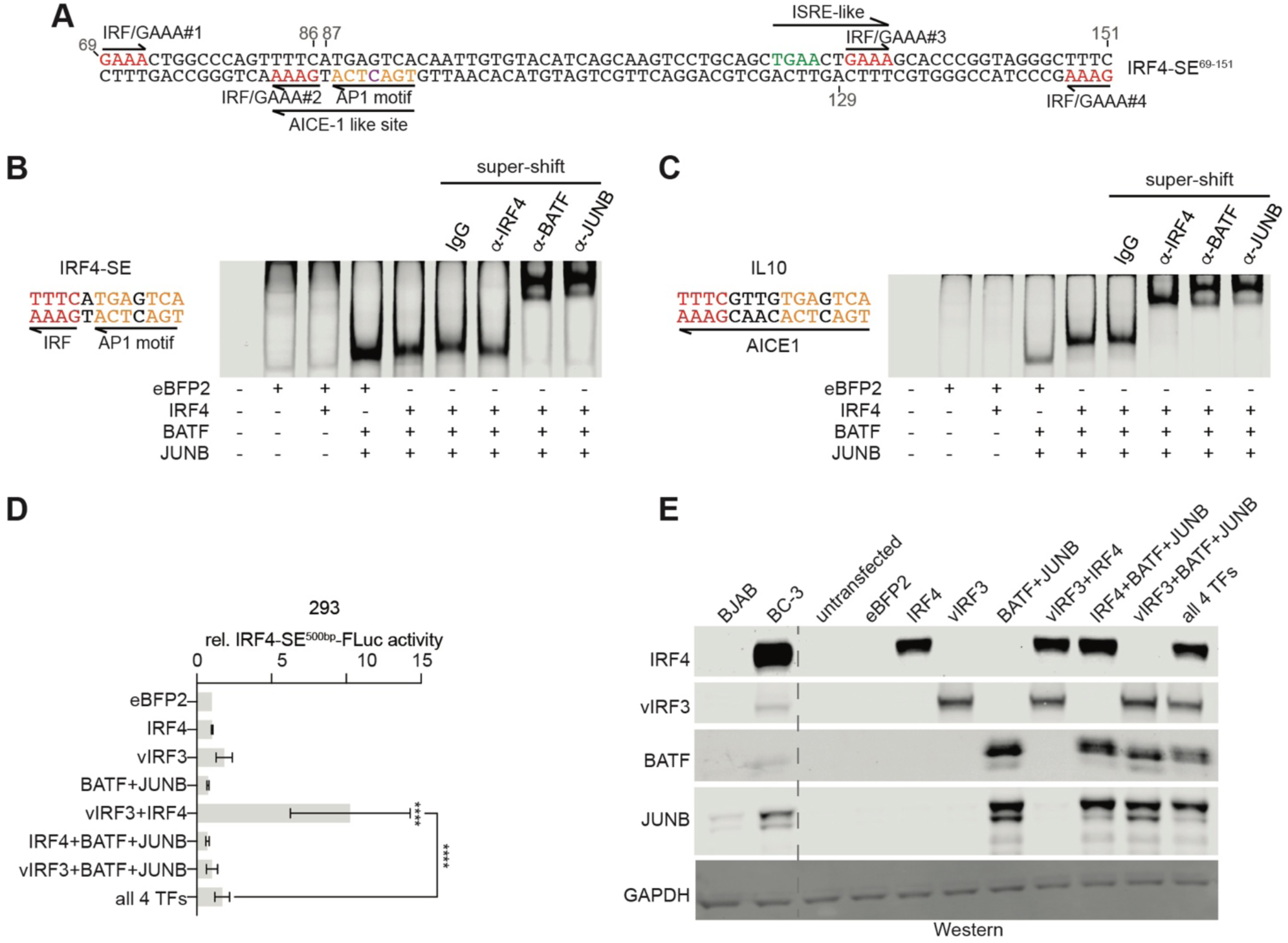
The minimal vIRF3/IRF-responsive sequence in the IRF4-SE contains a functional AP1 site, but not an AICE motif. **(A)** Sequence of nts 69-151 of the IRF4-SE^500bp^ element. Candidates for known motifs are in color. Arrows mark the orientation of each motif. **(B)** Electrophoretic Mobility Shift Assays (EMSAs) using a probe containing an AICE1-like motif from the IRF-SE4 and nuclear extracts from transfected 293T cells expressing combinations of EBFP2, IRF4, BATF, and JUNB as indicated. The AP1 (orange) and IRF (red) motifs are in color. Supershift assays were performed using antibodies listed above the panel. Representative of 2 independent repeats. **(C)** EMSA as in panel B but using a positive control probe containing the canonical AICE1 site from IL10. Representative of 2 independent repeats. **(D)** Results from dual-luciferase reporter assays as described in Fig. 1C, except that different combinations of BATF and JUNB were included. Error bars represent SD from 3 biological replicates. ****, *adj. p <* 0.001 from One-Way ANOVA followed by Tukey’s post hoc tests. **(E)** Western blot analyses of 293 cells transfected with the indicated expression vectors under conditions used in panel (D). GAPDH served as a loading control. BC-3 is a control for expression levels in PEL cells. BJAB is a control for expression levels in a KSHV-negative B cell lymphoma cell line. Representative of 2 independent repeats.

Electrophoretic mobility shift assays (EMSAs) showed that the IRF4-SE AICE-1-like motif can bind to the BATF-JUNB AP1 complex but fails to subsequently recruit IRF4 (Fig. 2B). We chose JUNB for this assay, since it is more highly expressed in PEL cells than other JUN family members (8, and Fig. S3). In contrast, we obtained IRF4 recruitment upon BATF-JUNB AP1 complex binding by a positive control AICE1 motif from IL10 (Fig. 2C), as reported (34). In our reporter assay, the additional co-transfection with BATF and JUNB did not increase IRF4-SE^500bp^ reporter activity, suggesting that the role of vIRF3 and IRF4 on this element may, surprisingly, be independent of BATF and JUN. BATF and JUNB instead inhibited vIRF3/IRF4-mediated IRF4-SE element activation (Fig. 2D-E), potentially pointing towards competition with vIRF3.

### The DNA binding capacity of IRF4 is critical for cooperative IRF4-SE activation

The identified minimal vIRF3/IRF4-responsive element contains several IRF-related motifs, including a candidate ISRE-like site (Fig. 2A), suggesting that IRF4 directly binds these elements to activate reporter expression. We, therefore, used a published IRF-motif-binding-deficient mutant of IRF4 (IRF4^R98A/C99A^) (32,33,45) to ask whether the IRF4 DNA-binding capacity is important for cooperative reporter activation by this element. In DNA pulldown assays, a biotinylated fragment spanning the minimal vIRF3/IRF4-responsive region (IRF4-SE^69-151^) and an additional 15 bp on either side associated with wt but not R98A/C99A-mutant IRF4, confirming the DNA-binding deficiency of IRF4^R98A/C99A^ (Fig. 3A). The biotinylated IRF4-SE probe recovered IRF4 independently of vIRF3 co-transfection. In turn, the biotinylated probe recovered vIRF3 independently of whether it was transfected with wt or R98A/C99A-mutant IRF4. Interestingly, wt IRF4 and vIRF3 showed a lower degree of association with a randomized biotinylated DNA fragment (N6, Table S2) designed to lack any IRF/GAAA motifs or other known TF binding sites. This negative control sequence did not activate Fluc reporter expression after co-transfection with vIRF3 and IRF4 (Fig. 3B). We observed similar background association with other biotinylated negative control probes but only marginal recovery in the presence of unbiotinylated probes (see below for quantification). Co-immunoprecipitation of IRF4 and vIRF3 from nuclear extracts, as published (8), shows that the IRF4^R98A/C99A^ mutation does not affect association with vIRF3 (Fig. 3C). In reporter assays, the DNA-binding-deficient IRF4 mutant was unable to cooperate with vIRF3 to activate either IRF4-SE^500bp^ or the minimal IRF4-SE^69-151^ region (Fig. 3D-E). Interestingly, we observed significant regulation of the IRF4-SE^69-151^ by vIRF3 alone and the co-transfection of R98A/C99A-mutant IRF4 did not interfere with this activation by vIRF3 (Fig. 3E). Together, these results suggest that the ability of IRF4 to associate with DNA is critical for its cooperation with vIRF3 on the IRF4-SE element.

**Figure 3.**
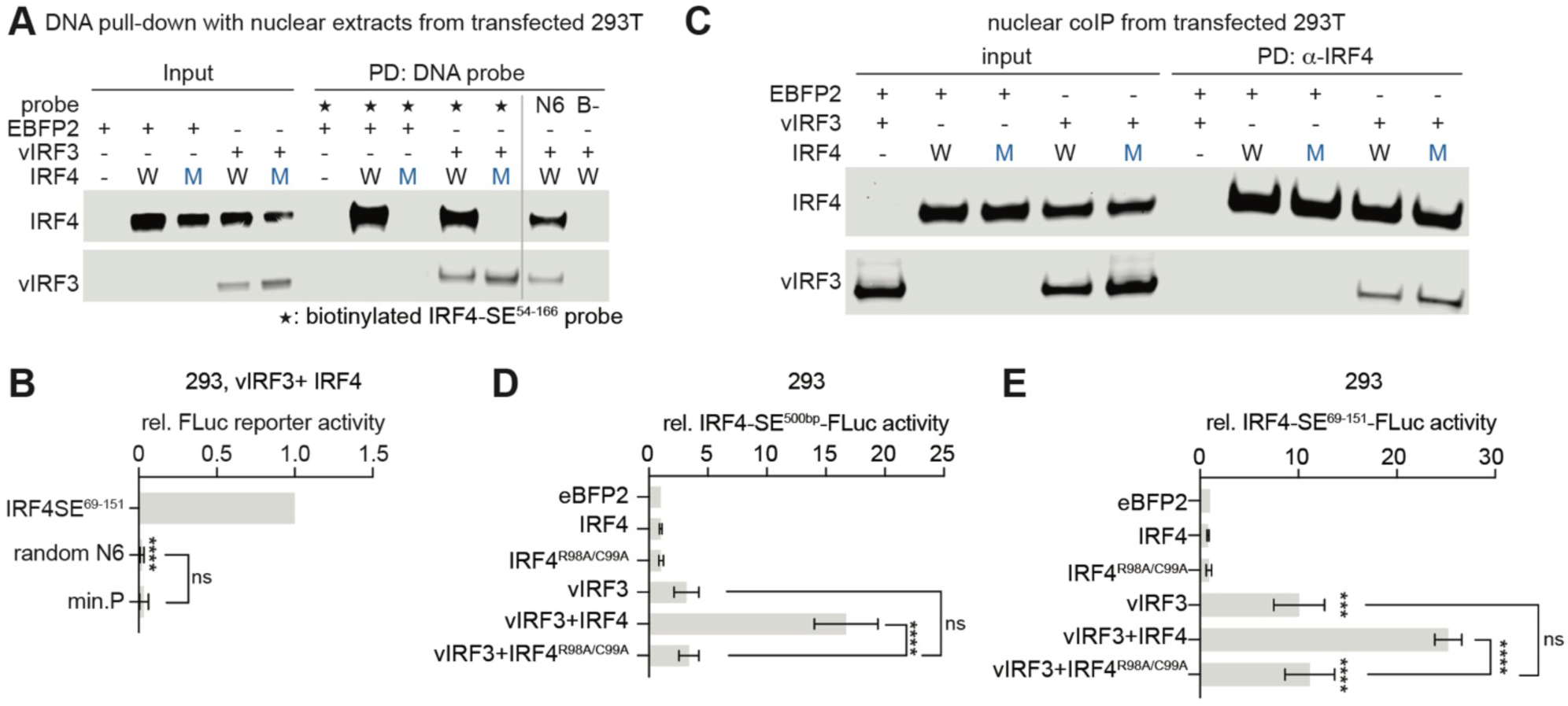
The DNA-binding capacity of IRF4 is required for its cooperation with vIRF3. **(A)** DNA pull-down assays using nuclear extracts from 293T transfected with the indicated protein expression vectors confirm that DNA binding deficient IRF4^R98A/C99A^ cannot associate with a biotinylated probe comprised of nts 54-166 of the IRF4-SE^500bp^ (IRF4-SE^54-166^, designated by “*”). N6 denotes a biotinylated randomized control probe (see Table. S2), B-denotes an unbiotinylated IRF4-SE^54-166^ probe. Recovered proteins were analyzed by Western blotting for IRF4 and vIRF3. Representative of 2 independent repeats. **(B)** Results from dual-luciferase reporter assays as described in Fig. 1D, showing that the N6 control sequence does not respond to co-transfection of vIRF3 and IRF4. IRF4-SE^69-151^ and minimal promoter (min. P) served as positive and negative controls, respectively. **(C)** Nuclear co-immunoprecipitation (Co-IP) with anti-IRF4 antibody, as published (8), but using nuclear lysates of 293T that were transfected with the indicated plasmids, shows that IRF4^R98A/C99A^ and IRF4^wt^ similarly associate with vIRF3. Co-precipitates were analyzed by Western blotting for IRF4 and vIRF3. Representative of 2 independent repeats. **(D)** Results from dual-luciferase reporter assays in 293 using the IRF4-SE^500bp^ reporter, as described in Fig. 1C, but including the IRF4^R98A/C99A^ mutant to compare the effect of IRF4^wt^ and _IRF4_R98A/C99A. **(E)** As in panel (D) but using the IRF4-SE^69-151^ reporter. Throughout, error bars represent SD from 3 biological replicates. ns, not significant; ***, *adj. p <* 0.001; ****, *adj. p <* 0.0001, calculated using One-Way ANOVA followed by Tukey’s post hoc tests.

### Separate sequences mediate IRF4-SE activation by vIRF3 and cooperation with IRF4

Given the importance of IRF4’s ability to bind to DNA for its cooperation with vIRF3, we directly tested the roles of the IRF/GAAA-related motifs and the AP1 motif in the context of the minimal IRF4-SE^69-151^ reporter. For this, we mutated each IRF/GAAA motif alone, the four IRF/GAAA motifs together, candidate composite sites (i.e., the AICE1-like and ISRE-like motifs), the AP1 motif, and the TGAA candidate ISRE half-site (Fig. 4A). Upon co-transfection of 293 with expression vectors encoding vIRF3 and IRF4, mutation of each individual IRF/GAAA motif led to a modest, but significant, reduction in reporter activity (Fig. 4B). Mutating the 4 IRF/GAAA motifs together strongly decreased IRF4-SE^69-151^ activity. Mutation of the AICE1-like-motif reduced reporter activity to background level. Mutation of the AP1 site (TGAnTCA) had a significantly stronger effect than eliminating the closely spaced IRF motif (IRF/GAAA #2) (*adj. p*=0.0047, one-way ANOVA with Tukey’s multiple comparisons test). In contrast, the strong reduction in reporter activity after mutating the entire AICE1-like motif or the AP1 site alone was not significantly different. Mutating the ISRE-like motif or either candidate half site (i.e., IRF/GAAA #3 or TGAA) strongly impaired reporter activity and any differences between these three mutants did not reach significance in this assay. Interestingly, only the AP1 and AICE1-like site mutations significantly reduced reporter activation by vIRF3 alone (Fig. 4C). These results suggest that the AP1 motif is critical for the response to vIRF3, while the IRF/GAAA-related motifs, including three apparently isolated IRF/GAAA motifs and an ISRE-like motif, are critical for the cooperation of vIRF3 with IRF4. Thus, the elements required for the basic response to vIRF3 and its cooperation with IRF4 are genetically separable. Importantly, the AP1 site and the IRF/GAAA and ISRE-like motifs do not form a known composite site but instead cooperate within a complex and extended composite element.

**Figure 4.**
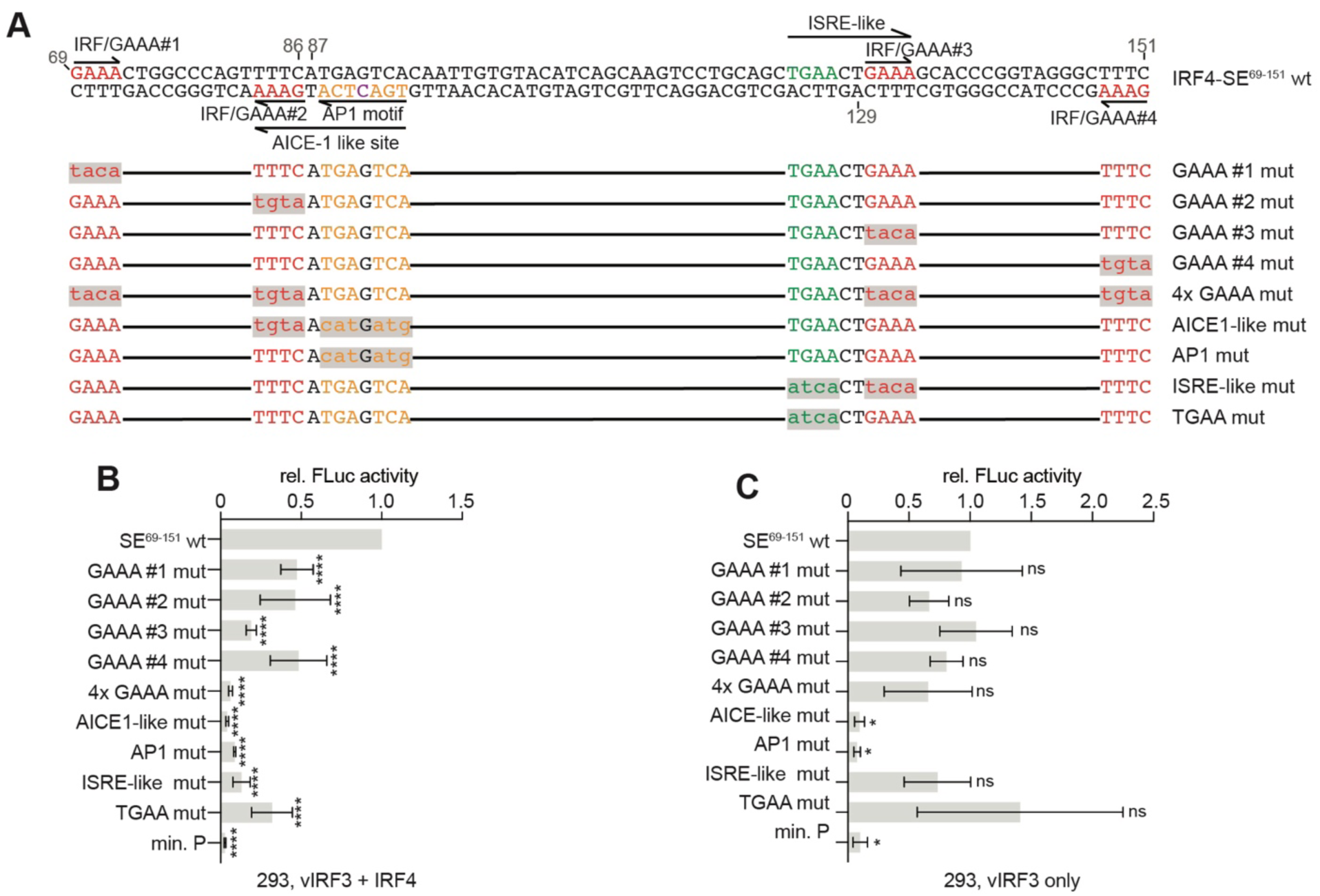
The vIRF3-responsiveness of the IRF4-SE depends on the AP1 site, while cooperation with IRF4 depends on IRF/GAAA motifs, including a candidate ISRE. **(A)** Sequences of wt IRF4-SE^69-151^ and mutants used in this figure. IRF/GAAA motifs are in red, the AP1 site is in orange, and the TGAA IRF-like half site of the candidate ISRE is in green. Mutated nts are in lowercase with grey shading. Unchanged nts are shown as black lines. **(B)** Dual-luciferase reporter assays showing relative wt and motif-mutant IRF4-SE^69-151^ reporter activities following co-transfection with vIRF3 and IRF4. Normalization to results obtained with EBFP2 was performed as in Fig. 1D and significance compared to the wt reporter was determined using One-Way ANOVA followed by Dunnett’s post hoc tests. Error bars represent SD from 3 biological replicates for bar graphs. ns, not significant; ****, *adj. p <* 0.0001. **(C)** Dual-luciferase reporter assays as described in panel B but testing transfection of vIRF3 only. Normalized as in panel B. Results from the AICE-like and AP1 mutants differed significantly from the wt IRF4-SE^69-151^, determined using One-Way ANOVA followed by Dunnett’s post hoc tests. Error bars represent SD from 3 biological replicates for bar graphs. ns, not significant; *, *adj. p <* 0.05.

### IRF/GAAA, ISRE-like, and AP1 motifs mediate IRF4-SE activity in PEL cells

We next confirmed the importance of the IRF/GAAA motifs, the ISRE-like motif, and the AICE1-like motif using lentiviral EGFP-based IRF4-SE^69-151^ reporters in the PEL cell line BC-3 (Fig. 5A-B), using the vector shown in Fig. 1G. While the minimal IRF4-SE^69-151^ element was sufficient to increase EGFP expression in BC-3, reporter activation was weaker than with the IRF4-SE^500bp^ element (compare Fig. 5A-B and Fig. 1H-I), not allowing for sufficient distinction of EGFP-expressing cells from untransduced cells. Notably, IRF4-SE^69-151^ showed improved activity over IRF4-SE^500bp^ in 293 cells. The reduced activity of IRF4-SE^69-151^ in BC-3 may reflect differences between the transfection- and transduction-based assays, the slightly different ratios of vIRF3 and IRF4 expression we achieve in 293 compared to those in BC-3 (Fig. 1B), the different cellular context, and positive regulatory roles for sequences outside the core element in BC-3 that are not captured in 293. It is furthermore possible that sequences outside nts 69-151 are subject to unphysiological negative regulation in 293. Since SEs are defined by their large size and contain densely packed regulatory elements, the presence of additional regulatory elements in the IRF4-SE is expected. Despite this, mutations in this construct confirm the importance for the AICE1-like and ISRE-like motifs in IRF4 reporter activation in PEL cells (Fig. 5A-B).

**Figure 5.**
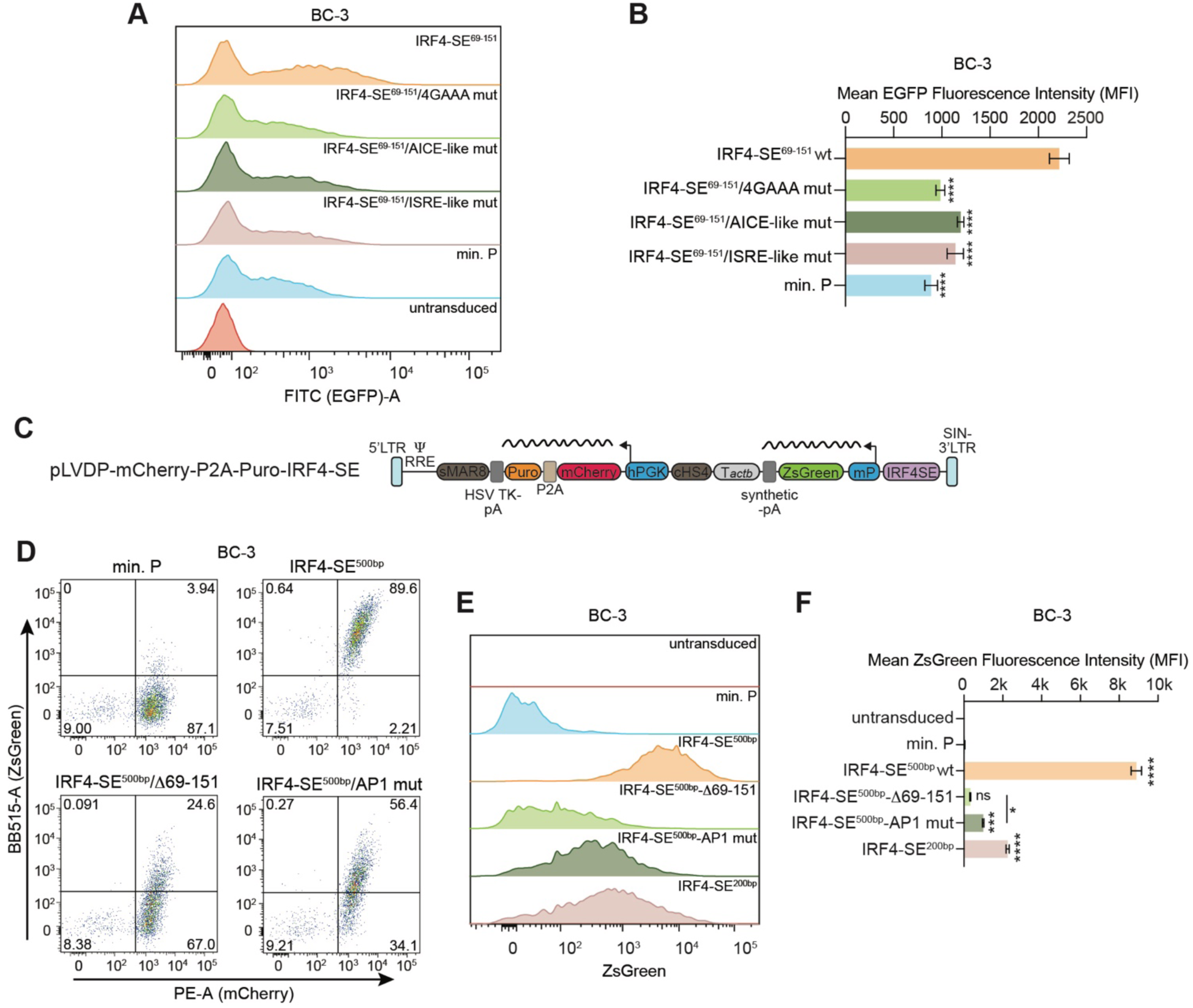
The AP-1 motif is critical for IRF4-SE reporter activity in the BC-3 PEL cell line. **(A)** Representative histogram comparing EGFP expression 3 days after transduction of BC-3 cells with pLS-mP lentiviral reporters containing the indicated wt or mutant IRF4-SE^69-151^ elements at equal MOI (∼1). Quantified over 3 repeats in panel (B). **(B)** Quantification of EGFP mean fluorescence intensity of EGFP-positive BC-3, defined based on an untransduced negative control, over 3 independent experiments as in panel (A). **(C)** Schematic of the dual-reporter lentiviral construct used in panels (**D-F**). Abbreviations in addition to those in Fig. 1G are scaffold/matrix attachment (S/MAR) region 8 (sMAR8), herpes simplex virus thymidine kinase polyadenylation signal (HSV TK pA), puromycin resistance cassette (Puro), P2A ribosomal skipping peptide, mCherry coding sequence (mCherry), and chicken hypersensitive site 4 (cHS4) from the β-globin locus. Tactb indicates a sequence facilitating efficient transcription termination of the upstream ZsGreen reporter cassette and downstream of a synthetic polyadenylation signal (pA). Wavy lines indicate expected transcripts after transduction. **(D)** Quadrant plots to illustrate mCherry and ZsGreen reporter expression in puromycin-selected BC-3 cells 3 days after transduction with the indicated pLVDP-mCherry-P2A-Puro-minP vectors at MOI 0.3. Gated for single, live cells. Numbers in quadrants indicate percentages of the parent population. Representative of 3 independent repeats, which are further analyzed and quantified in panels (E-F). **(E)** Representative histogram comparing ZsGreen reporter expression in transduced cells as in panel (D). Gated as in D and additionally for mCherry-positive cells. **(F)** Quantification of ZsGreen mean fluorescence intensity in transduced BC-3 cells over 3 independent experiments as in panel (E). Significance was compared to min.P, unless indicated by lines. ZsGreen expression from all constructs was furthermore significantly reduced compared to IRF4-SE^500bp^, *adj. p <* 0.0001, determined using One-Way ANOVA followed by Tukey’s post hoc tests. Throughout, error bars represent SD from 3 biological replicates for bar graphs. ns, not significant; *, *adj. p <* 0.05; **, *adj. p <* 0.01; ***, *adj. p <* 0.001; ****, *adj. p <* 0.0001 from One-Way ANOVA followed by Tukey’s post hoc tests.

To more readily distinguish transduced from untransduced cells and thereby increase the resolution of this assay, we developed a lentiviral vector that combines a promoter/enhancer element-driven ZsGreen reporter cassette with an insulated mCherry-P2A-Puro^R^ expression cassette (Fig. 5C), based on a reported design (46). Inclusion of the mCherry-P2A-Puro^R^ cassette enables convenient lentiviral titration and reporter analysis by selection with puromycin and/or gating for mCherry-expressing cells (Fig. 5D and S4). Using this approach, we showed that mutation of the AP1 site or deletion of the minimal 83 bp responsive element (nts 69-151) in the context of the IRF4^500bp^ reporter strongly impaired its activity in PEL cells (Fig. 5D-F), confirming the requirement for the AP1 site and surrounding sequences in BC-3 cells. These experiments furthermore directly confirmed that shortening the IRF4^500bp^ decreases its activity in BC-3 cells, even when including additional sequence context compared to IRF4-SE^69-151^, as we did in IRF4^200bp^, which includes the first 200bp of IRF4^500bp^ (Fig. 5E-F).

### The association of the IRF4-SE with vIRF3 and IRF4 is genetically separable

To explore the role of the identified motifs in recruiting IRF4 and vIRF3 to the minimal vIRF3/IRF4-responsive element, we generated biotinylated DNA fragments carrying a subset of the IRF- and AP1-related mutations. Control pulldowns with two biotinylated negative controls showed similar background association with IRF4 and vIRF3 in BC3 nuclear extracts (probes N2, N6, Figs. 6A-B, S4). The WT IRF4-SE fragment recovered 2-3-fold more IRF4 and vIRF3 compared to the negative control pulldowns, suggesting sequence-specific association of these proteins, despite background binding. Interestingly, mutation of the four GAAA motifs or the ISRE-like motif reduced the association of IRF4 to background levels. These results suggest that IRF4 is recruited specifically through GAAA motifs, and, most importantly, an ISRE-like element with a non-canonical TGAA 5’-half site. The 4GAAA or ISRE-like mutations did not reduce association with vIRF3 (Fig. 6C). Instead, the association with vIRF3 depended on the AICE1-like motif, consistent with the importance of the AP1 motif in the assays above. Mutation of the AICE1-like motif did not decrease IRF4 association, further confirming that this motif is unlikely to represent a functional AICE1 element. Together, these data suggest that the association of vIRF3 and IRF4 with the minimal IRF4-SE element is genetically separable, at least in the context of this *in vitro* assay using biotinylated DNA fragments.

**Figure 6.**
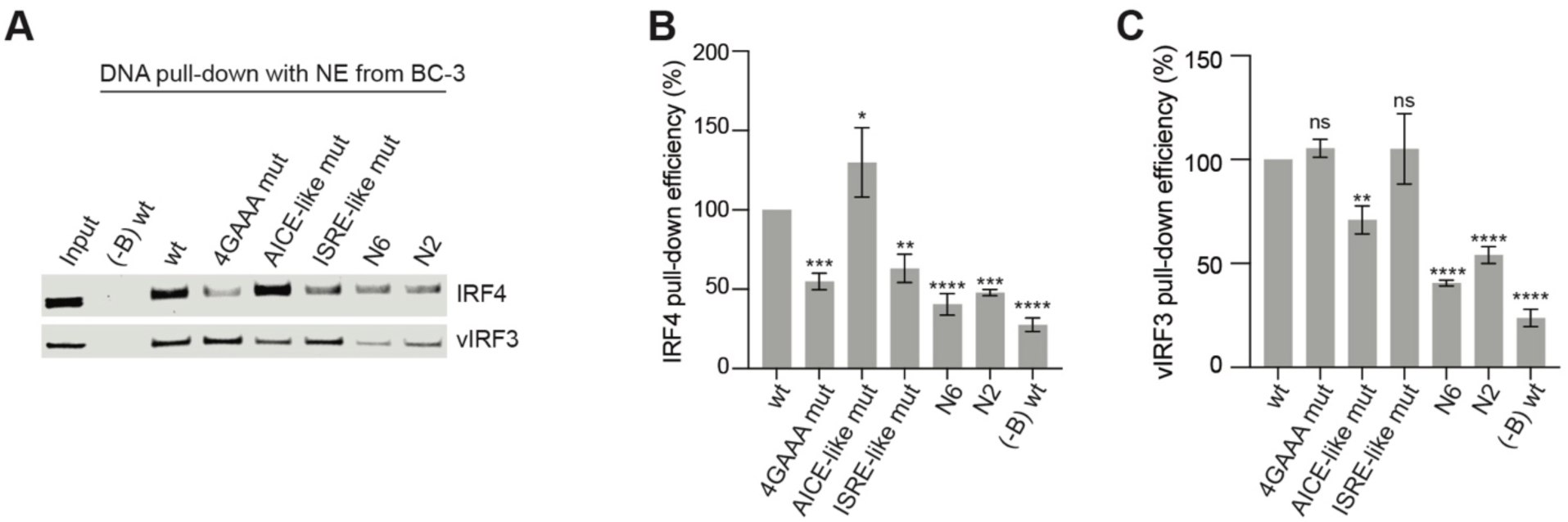
vIRF3 and IRF4 association with the vIRF3/IRF4-responsive elements depends on distinct motifs. **(A)** DNA pull-down assays, as in Fig. 3A, but using BC-3 nuclear extracts and a subset of the motif mutants from Fig. 4. Probes are biotinylated unless marked with (-B). N2/6 are randomized negative control probes (see Table S2). Representative of 3 independent repeats, which are quantified in panels (B-C). **(B-C)** Quantification of IRF4 (B) and vIRF3 (C) pull-down efficiency from three independent experiments as shown in (A). Error bars represent SD from 3 biological replicates. ns, not significant; *, *adj. p <* 0.05; **, *adj. p <* 0.01; ***, *adj. p <* 0.001; ****, *adj. p <* 0.0001 from One-Way ANOVA followed by Tukey’s post hoc tests.

### IRF and AP1-related motifs are insufficient for the vIRF3/IRF4 responsiveness of the IRF4-SE

Due to the critical role of the IRF, ISRE-like, and AP1 motifs, we next asked if these sites alone are sufficient to confer vIRF3/IRF4-responsiveness in an otherwise randomized context. We generated reporters with randomized sequences of all nts outside the motifs mentioned above (R1 and R2). We additionally generated reporters that preserved known motifs but randomized shorter sequences separating them (R3-R5, Fig. 7A). Of these reporters, only R4, which preserves sequences separating the AP1 and ISRE-like motifs, had detectable reporter activity in 293 cells, achieving ∼30% of wt reporter activation upon co-transfection of vIRF3 and IRF4 (Fig. 7B). Taken together with results from above, this result suggests that the IRF, ISRE-like and AP-1 motifs are critical for the activity of the minimal vIRF3/IRF4-responsive element but insufficient. In BC-3 cells, none of the partially randomized mutants substantially activated the lentiviral IRF4-SE^61-151bp^ EGFP reporter (Fig. 7C-D), confirming that the IRF- and AP1-related motifs are also insufficient in this context. The differential activity of R4 in 293 and BC-3 may be due to reporter integration status, and/or differences in cellular context, including the availability or stoichiometry of vIRF3, IRF4, and cofactors.

**Figure 7.**
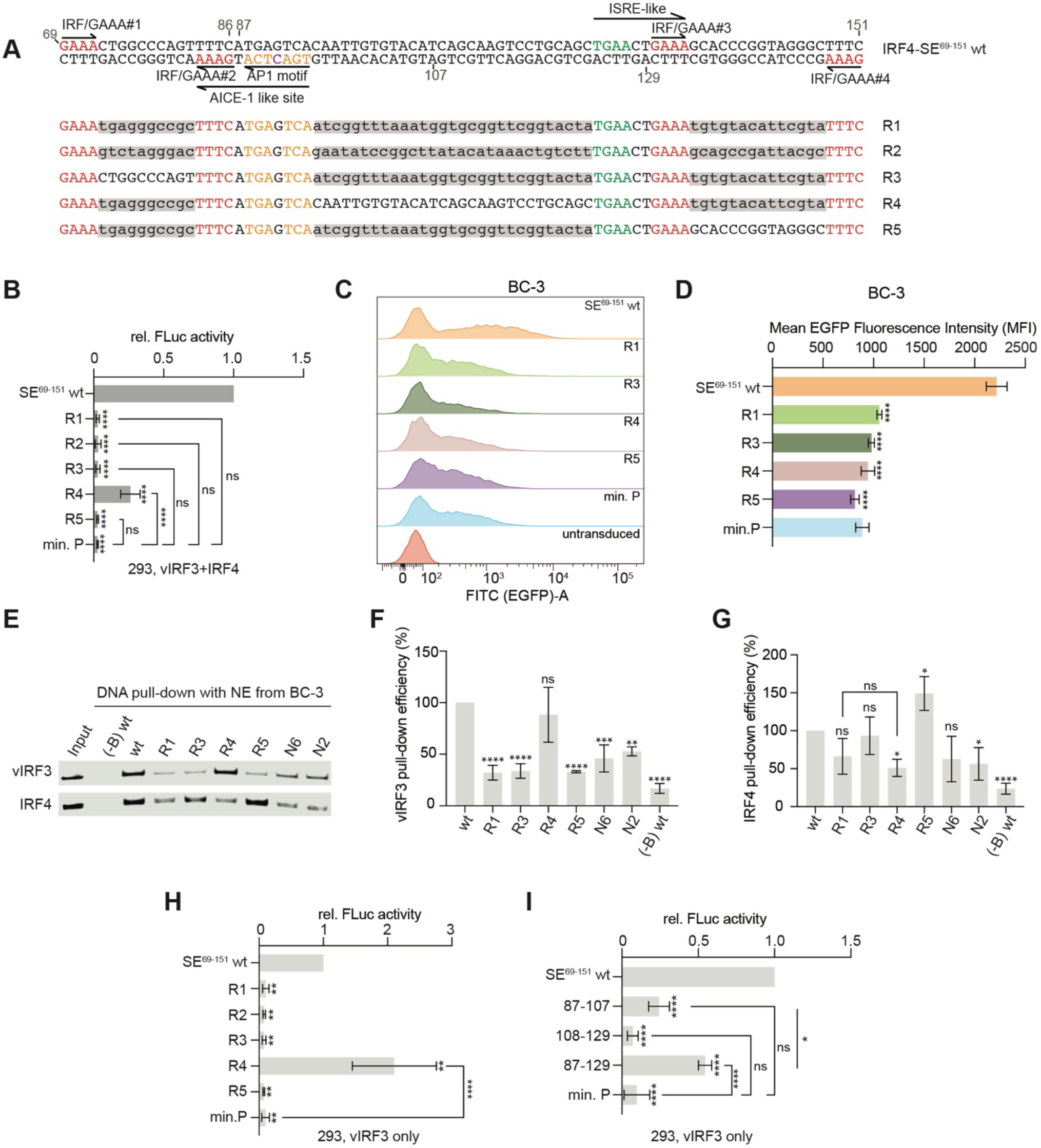
The identified motifs are insufficient for full vIRF3/IRF4 responsiveness of the IRF4-SE. **(A)** Schematic of IRF4-SE^69-151^ mutants with intact identified motifs and fully or partially randomized context (R1-5). Mutant sequences are in lowercase with grey shading. Previously identified motifs are in color and unchanged. **(B)** Dual-luciferase reporter assays showing relative wt and randomized-mutant IRF4-SE^69-151^ reporter activities following co-transfection with vIRF3 and IRF4. Normalization to results from EBFP2 and wt IRF4-SE^69-151^ was performed as in Fig. 1D. Significance is indicated relative to the wt vector. **(C)** Histogram comparing EGFP expression 3 days after transduction of BC-3 cells with pLS-mP lentiviral reporters containing the indicated wt or mutant IRF4-SE^69-151^ elements at equal MOI (∼1). Representative of 3 independent repeats. **(D)** Quantification of EGFP mean fluorescence intensity in EGFP-positive BC-3 cells, defined based on an untransduced negative control, over 3 independent experiments as in panel (C). **(E)** DNA pull-down assays, as above, using the indicated sequences and BC-3 nuclear extracts. Probes were biotinylated, except when marked (-B). N6 and N2 are randomized negative control probes as in Fig.6A and Table S2. Representative of 3 independent repeats, which are quantified in panels (F-G). **(F-G)** Quantification of vIRF3 (F) and IRF4 (G) pull-down efficiency from three independent experiments as shown in (E). **(H)** Dual-luciferase reporter assays showing relative wt and randomized-mutant IRF4-SE^69-151^ reporter activities following expression of vIRF3 only. Normalization to results from EBFP2 and the wt reporter was performed as in Fig. 1D. Significance is relative to the wt reporter, unless indicated for specific comparisons. **(I)** Dual-luciferase reporter assays showing relative wt and short IRF4-SE fragment reporter activities following transfection with vIRF3 only. Normalization to results from EBFP2 and the wt reporter was performed as in Fig. 1D. Significance is relative to the wt reporter, unless indicated for specific comparisons. Throughout, error bars represent SD from 3 biological replicates for bar graphs. ns, not significant; *, *adj. p <* 0.05; **, *adj. p <* 0.01; ***, *adj. p <* 0.001; ****, *adj. p <* 0.0001 from One-Way ANOVA followed by Tukey’s post hoc tests.

To further characterize the role of the sequences separating the IRF-related motifs, we subjected mutants R1-5, together with the N2 and N6 control fragments to DNA pulldown assays with BC-3 nuclear extracts. Sequences R1, R3, and R5 did not specifically associate with vIRF3, while mutant R4, which preserves the sequences separating the AP1 and ISRE-like motifs and was vIRF3-responsive in 293 reporter assays, showed wt levels of vIRF3 association (Fig. 7E-F). Interestingly, mutant R4 failed to specifically recruit IRF4 from BC-3 nuclear extracts, although it retains all IRF/GAAA-related motifs and the ISRE-like element, potentially explaining why this reporter was not active in BC-3 cells (Fig. 7E, G). These pulldown data suggest that the sequences separating the AP1 and ISRE-like sites promote recruitment of vIRF3 to the IRF4-SE. Accordingly, mutant R4 was fully responsive to vIRF3 in the context of the minimal IRF4-SE^69-151^ reporter (Fig. 7H). Furthermore, shorter fragments that retained the AP1 motif, but lacked any IRF motifs responded to vIRF3 (Fig. 7I). Interestingly, a fragment encompassing nts 87-129 of the IRF4-SE^500bp^ conferred significantly greater activation than a fragment including only nts 87 to 107, suggesting that the role of the sequences separating the AP1 site and the ISRE-like motif is not simply due to the immediate sequence context downstream of the AP1 site, but involves a relatively extended sequence. Together with the data above, these results confirm that recruitment of vIRF3 and IRF4 to the IRF4-SE are genetically separable and require not only IRF, ISRE-like, and AP1 motifs but also additional sequences.

### The PEL MYC-SEs are vIRF3/IRF4-responsive SEs

We finally sought to extend our analyses to other candidates for vIRF3/IRF4-responsive SEs. For this, we chose the 2 PEL MYC-SEs, located ∼500kb and 375kb downstream of the MYC transcription start site, respectively, due to their prominent occupancy by vIRF3 and IRF4 and the central importance of MYC for PEL cell survival (7,8,18,19,47). We inserted the SE sequences spanning the overlapping vIRF3/IRF4 ChIP-seq peaks into the transfected firefly reporter vector (Fig. S5). As for the IRF4-SE, vIRF3 and IRF4 cooperatively activated both MYC-SEs (Fig. 8A-B). Interestingly, the downstream MYC-SE^500kb^ element generated much higher reporter activity in 293 cells than the upstream MYC-SE^375kb^ element. While the MYC-SE^500kb^ reporter was furthermore substantially activated by vIRF3 alone, the MYC-SE^375kb^ reporter was not. Neither region was activated by IRF4 alone. 40bp deletion mutants mapped the region required for the vIRF3/IRF4-responsiveness of the stronger MYC-SE^500kb^ element to an ∼86 bp region between nts 121-207 (Fig. 8C), a region of similar size as the ∼83 bp vIRF3/IRF4-responsive region within the IRF4-SE. This region contains a canonical AICE2 motif near its 5’ end (Fig. 8D), which we confirmed as a functional AICE2 motif using EMSAs (Fig. S6A-B). Mutating the AICE2 motif greatly reduced reporter activation after co-transfection of vIRF3 and IRF4, while AP1 and IRF half-site mutants had more subtle, but significant effects (Fig. 8E). In contrast, the response to vIRF3 depended as much on the AP1 half site as the entire AICE2 motif, while mutating the IRF half site had no effect (Fig. 8F). These results suggest that vIRF3 requires the AP1 half site of the AICE2 motif, while cooperation with IRF4 may depend on the IRF half-site. Like for the IRF4-SE, co-transfection of JUNB and BATF did not enhance the response of the MYC-SE^500kb^ element to either vIRF3 alone or vIRF3 and IRF4 together, arguing against a recruitment of vIRF3 by this AP1 complex (Fig. S6C). Despite a functional AICE2 motif, co-transfection of IRF4, BATF, and JUNB was furthermore not sufficient to activate the MYC-SE^500kb^ reporter in 293 cells (Fig. S6C). In contrast to the more active MYC-SE^500kb^ element, the less active MYC-SE^375kb^ reporter does not contain a canonical AICE motif. Mutating a candidate AP1 site within MYC-SE^375kb^ furthermore did not affect reporter activity after IRF4 and vIRF3 co-transfection (Fig. S5, Fig. 8G), however, we note that this reporter contains additional motifs that match the AP1 consensus less stringently (see Fig. S5B for the sequence of this element). Using the dual lentiviral reporter vector (Fig. 5C), we finally confirmed the requirement for the AP1 site in the PEL cell line BC-3 (Fig. 8H-I). As for the IRF4-SE, shortening the reporter to a fragment spanning only nucleotides 81-247 of the original vector reduced its activity, showing that additional positive regulatory sequences exist in the originally tested 327 bp MYC-SE^500kb^ enhancer element.

**Figure 8.**
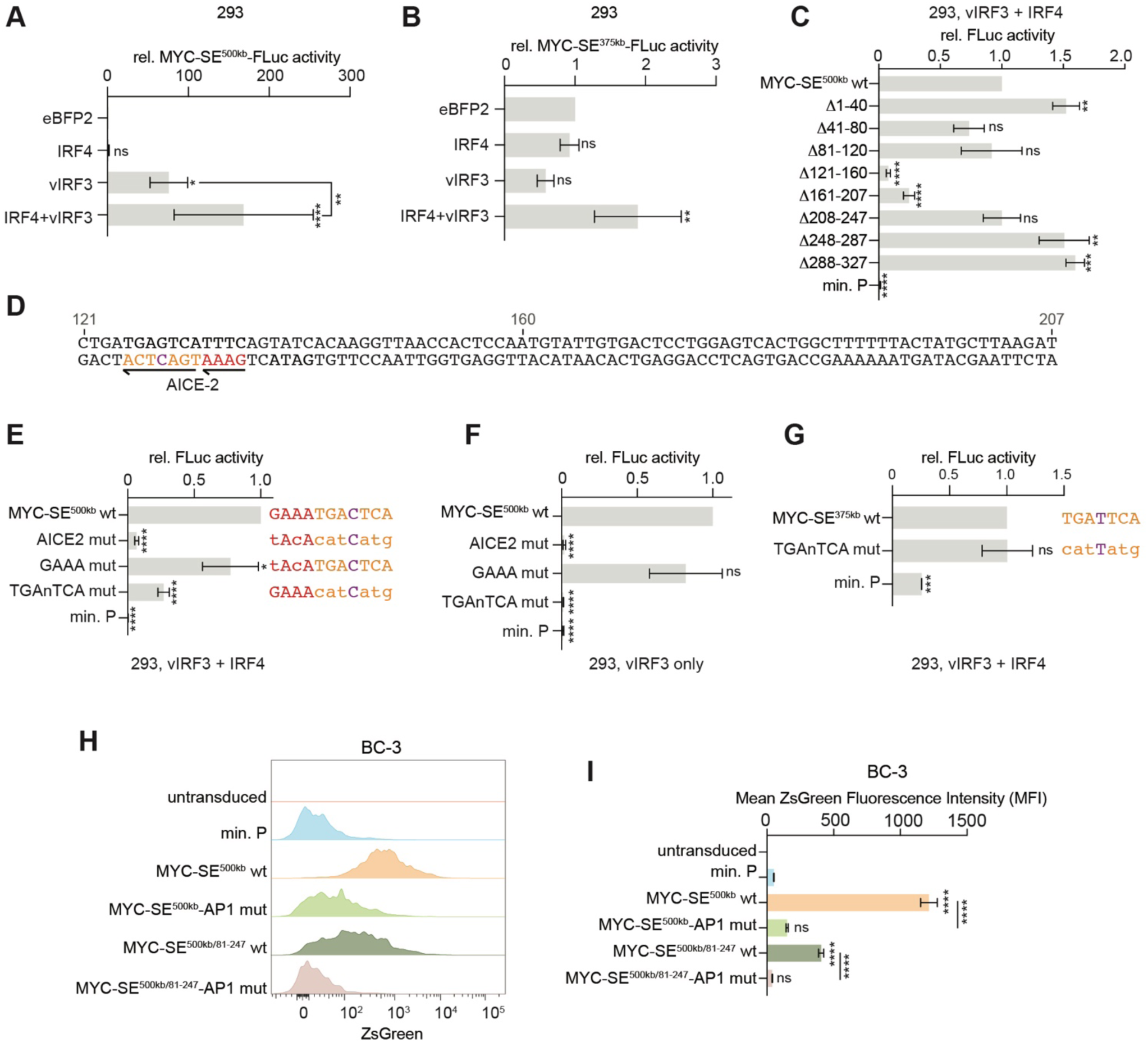
vIRF3/IRF4-responsiveness of the two PEL MYC-Ses. **(A)** Dual-luciferase reporter assays showing the relative activity of an FLuc reporter containing a 327 bp element centered on the vIRF3/IRF4 Chip-seq peaks in the MYC-SE located ∼500 kbp downstream of the MYC transcription start site (MYC-SE^500kb^, sequence in Fig. S6A), after co-transfection with expression vectors for the indicated proteins into 293. Results were normalized as in Fig. 1C, i.e., first to results from the RSV-LTR-RLuc reporter and then to results from EBFP2, which were set to 1. Results are from 7 independent repeats. Significance is relative to the EBFP2 control, unless indicated by lines. **(B)** As in panel A but using a reporter containing a 299 bp fragment centered on the vIRF3/IRF4 Chip-seq peaks in the MYC-SE located ∼375 kbp downstream of the MYC transcription start site (MYC-SE^375kb^, sequence in Fig. S6B). Data were normalized as in panel A. Results are from 5 independent repeats. Significance is relative to the EBFP2 control. **(C)** Results from dual-luciferase reporter assays showing wt or mutant MYC-SE^500kb^ activities after expression of vIRF3 and IRF4. Results from vIRF3/IRF4 co-transfection were normalized as in Fig. 1D, i.e. to results from the internal RSV-LTR-RLuc control, EBFP2, and the wt construct, which was set to 1. Results are from 3 independent repeats. Significance is relative to the wt reporter. **(D)** Sequence of MYC-SE^500kb^ nts 121-207. The candidate AICE2 site is marked in color. **(E)** Dual-luciferase reporter assays as in panel C but using wt or AICE2 site mutant MYC-SE^500kb^. Wt and mutant AICE2 motif sequences are next to each bar. Results are from 4 independent repeats. Significance is relative to the wt reporter. **(F)** Dual-luciferase reporter assays as in panel E but after expression of vIRF3 or EBFP2 only. Results are from 4 independent repeats. Significance is relative to the wt reporter. **(G)** Dual-luciferase reporter assays as in panel E but using wt and mutant MYC-SE^375kb^ reporters. The sequence of the wt or mutant AP1 motif is next to each bar. Results are from 3 independent repeats. Significance is relative to the wt reporter. **(H)** Histogram comparing ZsGreen reporter expression in BC-3 transduced with wt and mutant pLVDP-mCherry-P2A-Puro-MYC-SE^500kb^. Gated for single, live, mCherry-positive cells. Representative of 3 independent repeats, which are quantified in panel (I). **(I)** Quantification of ZsGreen mean fluorescence intensity in mCherry-positive BC-3 cells over 3 independent experiments as in panel H. Significance is relative to min. P, unless indicated by lines. Data were acquired together with Fig. 5D-F, and these experiments therefore share the untransduced and min. P controls. Throughout, error bars represent SD from 3 biological replicates for bar graphs. ns, not significant; *, *adj. p <* 0.05; **, *adj. p <* 0.01; ***, *adj. p <* 0.001; ****, *adj. p <* 0.0001 from One-Way ANOVA followed by Tukey’s post hoc tests.

## Discussion

The latent KSHV protein vIRF3 is essential for the survival of PEL cells (39), in part by promoting the SE-mediated overexpression of IRF4 and MYC (8). This study reveals that vIRF3 and IRF4 cooperatively activate both the IRF4-SE and a distal MYC-SE^500kb^, and that these elements rely on unusually long DNA sequences for responsiveness, approximately 80-90 bp, suggesting the presence of complex, composite regulatory sites.

Notably, IRF4 alone, or with BATF and JUNB, failed to activate reporter constructs, whereas vIRF3 alone could drive activity. IRF4 contributes to activity at both SEs but requires cooperation with vIRF3 for full activation. Our data show that the DNA sequences required for vIRF3 recruitment are genetically separable from those mediating its cooperation with IRF4, strongly arguing against recruitment of vIRF3 by IRF4. vIRF3 acts through critical AP1 motifs at both sites. In the case of the MYC-SE^500kb^, the AP1 motif forms a canonical AICE2 element with an adjacent IRF/GAAA motif. In the case of the IRF4-SE, the AP1 site did not participate in canonical AICE motifs but participated in non-canonical cooperation with an ISRE-like element and isolated IRF/GAAA motifs. These results highlight the essential role of vIRF3 in PEL-SE engagement and suggest that vIRF3 enables IRF4 to function in configurations that diverge from known composite binding motifs. A second MYC-SE, MYC-SE^375kb^, was not activated by vIRF3 alone and was significantly, but weakly, activated by co-transfection of vIRF3 and IRF4. This element may use an AP-1-independent mode of vIRF3 recruitment or vIRF3 might be recruited through a less optimal AP1 site that remains to be identified.

For the IRF4-SE, vIRF3 recruitment was not supported by sequences containing the AP1, IRF/GAAA, and ISRE-like motifs in randomized sequence contexts, and our data indicate that extended AP1-flanking sequences contribute to vIRF3 association and function. These results point to complex sequence requirements for vIRF3 recruitment and may help explain why our prior ChIP-seq analysis revealed only few high-confidence vIRF3 peaks (8). All ∼60 SE-associated vIRF3 peaks overlap IRF4 peaks, suggesting that vIRF3 specifically associates with regions that are enriched in IRF4 binding sites and recruit IRF4 in other settings. Accordingly, these sites overlapped IRF4-occupied SEs in other IRF4-dependent settings (8). While we cannot exclude that the low number of vIRF3 peaks is related to the performance of available vIRF3 antibodies, our data above favor a model in which vIRF3 targets a restricted set of SEs through multifactorial and context-dependent interactions, in cooperation with IRF4.

Although BATF/JUNB are established IRF4 partners, they do not appear to mediate vIRF3 activity. BATF and other BATF family members are only minimally expressed in 293 cells (Fig. S3), and BATF/JUNB co-transfection failed to enhance SE activity. Moreover, BATF knockout in PEL cells caused a slower loss of viability than the more rapid culture collapse following IRF4 or vIRF3 inactivation (8). Since ChIP-seq data indicate overlapping BATF co-occupancy at vIRF3-occupied SEs (8), we hypothesize that vIRF3 may functionally substitute for BATF/JUN at a small number of loci with specific sequence configurations, acting as a viral mimic of AP1-family IRF4 co-factors. This model is supported by reduced reporter activity upon BATF/JUNB co-transfection, pointing to competition, and by the observed AP1 motif dependence of for vIRF3 recruitment and function.

Our mutagenesis of the IRF4-SE further revealed that isolated IRF motifs, outside of canonical composite elements, can contribute to cooperative activation by vIRF3 and IRF4. Although IRF4 cannot efficiently bind such motifs alone, our findings suggest it may do so in the presence of vIRF3, or that vIRF3 modulates IRF4 binding behavior. While DNA pulldown assays did not show enhanced IRF4 recruitment in the presence of vIRF3, these assays may not capture the chromatin-level dynamics in PEL cells. Follow-up studies should therefore assess the influence of vIRF3 on IRF4 occupancy in native chromatin.

Because IRF4 or vIRF3 knockdown rapidly compromises PEL cell viability (7–9,39), we relied on ectopic expression experiments to isolate their roles. These approaches have inherent limitations, such as imperfect stoichiometry, absence of B-cell or PEL-specific chromatin features, and cell-type specific expression of other transcription factors. However, we validated key findings in PEL cells using reporter assays and DNA pulldowns, confirming that the mapped AP1 and IRF motifs are required for activity and recruitment. There are several important questions that should be addressed in future work. Specifically, the molecular basis for vIRF3 recruitment to DNA remains unresolved. The N-terminal region of vIRF3 folds similarly to the IRF DBD but lacks the conserved residues involved in recognition of the IRF/GAAA motif by IRFs. While vIRF3 is therefore not thought to bind to DNA directly, non-canonical binding to complex motifs may have been missed. Similarly, it is unknown how vIRF3 associates with IRF4 and whether this association is direct, although our results suggest that association of vIRF3 and IRF4 is independent of IRF4’s ability to bind to DNA.

In summary, our study defines sequence features underlying vIRF3-mediated SE activation in PEL cells and contributes to our understanding of how this KSHV protein co-opts host transcriptional machinery to sustain the oncogenic expression program in PEL. We hypothesize that vIRF3 may act as a functional analog of BATF/JUN AP1 complexes at specific loci, cooperating with IRF4 to activate SEs with non-canonical IRF4 binding motif architectures. These insights into viral manipulation of host gene regulation provide a starting point for dissecting SE biology in this and other IRF4-dependent contexts.

## Data Availability

No new data were generated in addition to those presented in the figures and tables. All vectors are available on request from EG.

## Funding

This work was supported by the National Cancer Institute (NCI) of the National Institutes of Health (NIH) under grant numbers R01-CA247619 (to EG) and R50-CA221848 (to ETB). The content is solely the responsibility of the authors and does not necessarily represent the official views of the National Institutes of Health.

## Conflict of interest disclosure

The authors declare that they have no competing interests.

## Author Contributions

Conceptualization: ZL, HKM, and EG. Cloning and investigation: ZL, HKM, CBM, MEH, JL. Formal analysis: ZL, ETB, EG. Visualization: ZL and EG. Supervision and validation: EG. Writing original draft: ZL and EG. Writing – review and editing: ZL and EG with feedback from all authors. Funding acquisition and project administration: ETB and EG.

## Acknowledgments

ZL, HKM, and JL were enrolled in the Northwestern University Master of Biotechnology program for part of this study. We would like to thank Raymond A. Xiong and Jesus A. Ortega for contributing vectors and protocols, respectively.

## Notes

### Competing Interest Statement

The authors have declared no competing interest.

