## Supplemental Figures 1-6 for "KSHV-encoded vIRF3 Cooperates with Cellular IRF4 to Drive Super-Enhancer Activity through Complex DNA Elements"

Liang et al.

This document contains Figures S1-6

See excel file for:
**Table S1** Sequences of all oligonucleotides or DNA fragments used for cloning
**Table S2** Oligonucleotides and PCR products used to generate DNA pulldown probes.
**Table S3** Oligonucleotides used for EMSA probes.
**Table S4** Antibodies and control IgG.

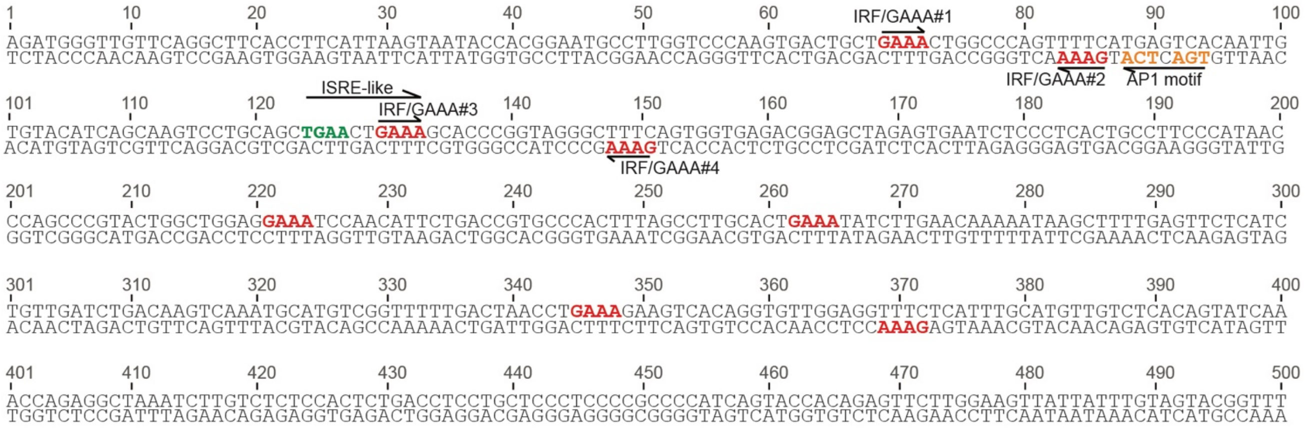


**Figure S1**
Sequence of IRF4-SE$^{500bp}$, corresponding to GRCh38/hg38 chr6:328,142-328,641. Candidates for known motifs are in color. Arrows mark the orientation of each motif.

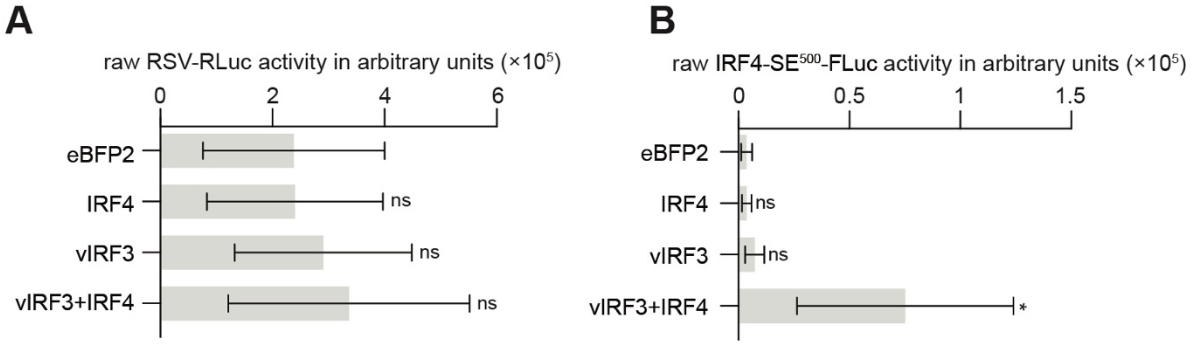


**Figure S2**
Raw RSV-RLuc (A) and IRF4-SE-FLuc activity (B) from the experiments shown in Fig. 1C.

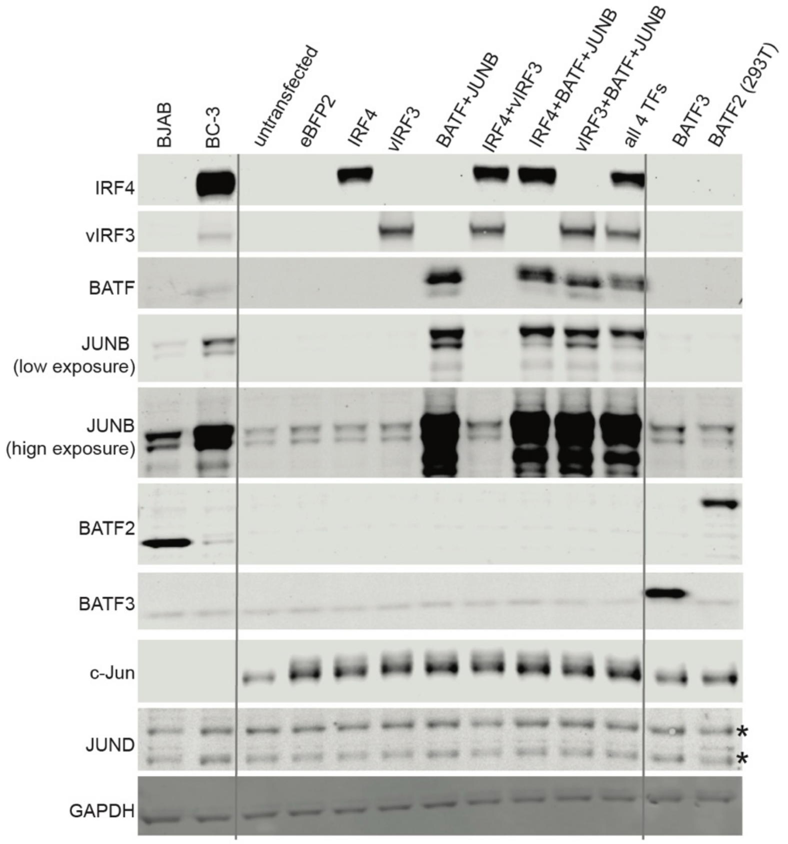


**Figure S3**
Extended version of the Western analyses shown in Fig. 2E, including an analysis of the endogenous expression levels of BATF and JUNB family members in 293 cells. BATF3 and BATF2 positive controls in the two rightmost lanes were derived from transfected 293 and 293T, respectively (constructs described in Methods). BJAB served as a KSHV-negative controls, BC-3 is a control PEL cell line. Grey lines separate controls, which were run on the same gel. These results suggest that 293 cells do not express high levels of BATF family members. Asterisks mark JUND bands we confirmed using siRNA-mediated knock-down experiments (not shown). Representative of n=2 repeats.

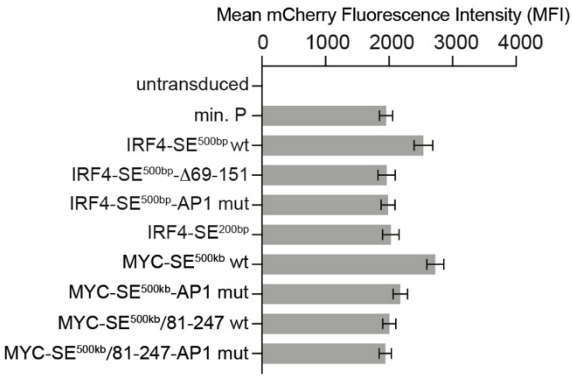


**Figure S4**
Quantification of mCherry mean fluorescence intensity in transduced BC-3 cells in the experiments shown in Fig. 5F and 8I.

**A** MYC-SE$^{500kb}$ (327bp)

```
1        10        20        30        40        50        60        70        80        90        100
ACAGCAAACCAAGAACCTGTGCAGCTGACACATATTGGGCACTTTCAGGAACACTAGCAGCATGAACACAGTGACTGAAGATTGAAACTAAGGAAACTAA
TGTCGTTTGGTTCTTGGACACGTCGACTGTGTATAACCCGTGAAAGTCCTTGTGATCGTCGTACTTGTGTCACTGACTTCTAACTTTGATTCCTTTGATT

101      110       120       130       140       150       160       170       180       190       200
TAGTAGTAAAGTCAGGGACTCTGATGAGTCATTTCAGTATCACAAGGTTAACCACTCCAATGTATTGTGACTCCTGGAGTCACTGGCTTTTTTACTATGC
ATCATCATTTCAGTCCCTGAGACTACTCAGTAAAGTCATAGTGTTCCAATTGGTGAGGTTACATAACACTGAGGACCTCAGTGACCGAAAAAATGATACG
                        AICE2

201      210       220       230       240       250       260       270       280       290       300
TTAAGATGTAACAAACTTATCTACCAGTTGAAAACCAGTTACATTTCTCCACTGCCTGCCACATAACATCAATCACTATTATAAAGAAGGATGGAAGGAA
AATTCTACATTGTTTGAATAGATGGTCAACTTTTGGTCAATGTAAAGAGGTGACGGACGGTGTATTGTAGTTAGTGATAATATTTCTTCCTACCTTCCTT

301      310       320      327
TTTAGTAAAAGGTGGTCCCCAGACTGT
AAATCATTTTCCACCAGGGGTCTGACA
```

**B** MYC-SE$^{375kb}$ (299bp)

```
1        10        20        30        40        50        60        70        80        90        100
CTTCATCTTAGAGGTGCAGGTGACAGTTGATAAGTACCTTGAAAGGCCTCCATGTGTGTGCAGGGCATTCTTTCGTAAGTTTCATCTGTACCATGTTAAG
GAAGTAGAATCTCCACGTCCACTGTCAACTATTCATGGAACTTTCCGGAGGTACACACACGTCCCGTAAGAAAGCATTCAAAGTAGACATGGTACAATTC

101      110       120       130       140       150       160       170       180       190       200
TGATTCAGGGCCACTATCTCTGAATTTGAAGAACCCTCAAGCTACTGGTTCTCCAGGACCTAAACCCTGGACTCCAGGCCACAGACAAGCTCTGAGTGAAA
ACTAAGTCCCGGTGATAGAGACTTAAACTTCTTGGGAGTTCGATGACCAAGAGGTCCTGGATTTGGGACCTGAGGTCCGGTGTCTGTTCGAGACTCACTTT

202     210       220       230       240       250       260       270       280       290       299
AATGAATCCTTCTTGCCGCTCCTCCCCTATGCTTCACATTGCTCCAAACCTACCTGCCTCCAACACGTGGTGCAGTTTCTACCTTCCCTTCCTTTGCA
TTACTTAGGAAGAACGGCGAGGAGGGGATACGAAGTGTAACGAGGTTTGGATGGACGGAGGTTGTGCACCACGTCAAAGATGGAAGGGAAGGAAACGT
```

**Figure S5**

**(A)** Sequences of the 327 bp MYC-SE$^{500kb}$ element used in Fig. 8A, corresponding to GRCh38/hg38 chr8:129,252,674-129,253,000. Relevant motifs are in color. The arrow marks the orientation of the AICE2 motif.

**(B)** Sequences of the 299 bp MYC-SE$^{375kb}$ element used in Fig. 8B, corresponding to GRCh38/hg38 chr8:129,124,349-129,124,647. Relevant motifs are in color. The arrow marks the orientation of the AICE2 motif.

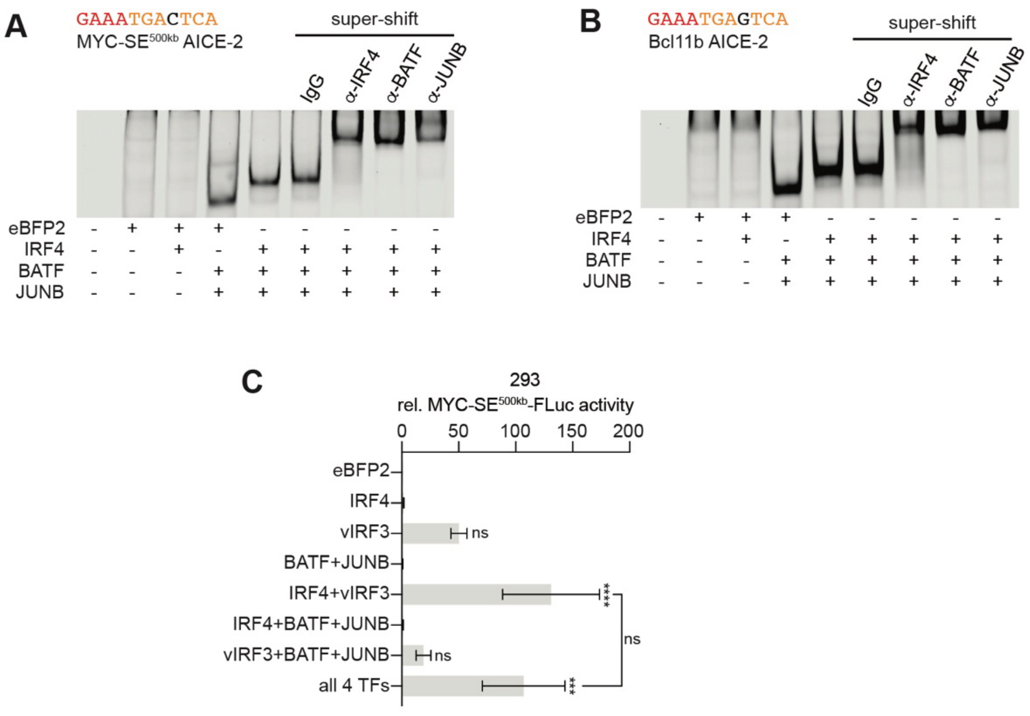


**Figure S6**

**(A-B)** EMSAs using nuclear extracts from transfected 293T cells expressing IRF4, BATF, and JUNB as for Fig 2A-B, and probes containing the AICE2 site from MYC-SE$^{500kb}$ (A) or the validated AICE2 site from Bcl11b (34) (B), including flanking sequences on both sides (see supplementary Table S3). The AP1 (orange) and IRF (red) half sites are in color. Supershift assays were performed using antibodies listed above the panels. Representative of n=2.

**(C)** Results from dual-luciferase reporter assays as described in Fig. 2D but using the MYC-SE$^{500kb}$ reporter. Error bars represent SD from 3 biological replicates. Significance was determined relative to the EBFP2 control, ns, not significant; ***, *adj.* $p < 0.001$; ****, *adj.* $p < 0.0001$ from One-Way ANOVA followed by Tukey's post hoc tests. Reporter activation by vIRF3 alone was significant using One-Way ANOVA followed by Dunnett's post hoc tests (*adj.* $p = 0.0428$).
